# Predictomes: A classifier-curated database of AlphaFold-modeled protein-protein interactions

**DOI:** 10.1101/2024.04.09.588596

**Authors:** Ernst W. Schmid, Johannes C. Walter

## Abstract

Protein-protein interactions (PPIs) are ubiquitous in biology, yet a comprehensive structural characterization of the PPIs underlying biochemical processes is lacking. Although AlphaFold-Multimer (AF-M) has the potential to fill this knowledge gap, standard AF-M confidence metrics do not reliably separate relevant PPIs from an abundance of false positive predictions. To address this limitation, we used machine learning on well curated datasets to train a <u>S</u>tructure <u>P</u>rediction and <u>O</u>mics informed <u>C</u>lassifier called SPOC that shows excellent performance in separating true and false PPIs, including in proteome-wide screens. We applied SPOC to an all-by-all matrix of nearly 300 human genome maintenance proteins, generating ∼40,000 predictions that can be viewed at predictomes.org, where users can also score their own predictions with SPOC. High confidence PPIs discovered using our approach suggest novel hypotheses in genome maintenance. Our results provide a framework for interpreting large scale AF-M screens and help lay the foundation for a proteome-wide structural interactome.

## Introduction

Most biological processes depend on the interaction of multiple proteins^1^. Stable protein-protein interactions (PPIs) form large cellular structures (e.g. the nuclear pore) and stable molecular machines (e.g. RNA polymerase), whereas transient interactions underlie dynamic processes ranging from signaling to DNA replication. The ∼20,000 proteins encoded by the human genome can theoretically combine in ∼200 million binary combinations, but current estimates suggest that only ∼1.5 million pairings represent functional PPIs^2^. Of these, only 50,000 (3%) have been identified^2^, and ∼8,000 (0.5%) are structurally resolved. These estimates, though necessarily imprecise, indicate that the vast majority of PPIs are both unknown and structurally inaccessible. Recognizing this massive knowledge gap, investigators have for decades sought to discover PPIs at scale ^3^. Experimental approaches toward this goal include yeast two hybrid assays^2,4^, as well co-immunoprecipitation^5^, column chromatography based complex fractionation^6^, and cross-linking coupled with mass spectrometry^7^. Computational approaches involve homology modeling^8^ and rigid body docking^9^. While these various methods have uncovered many new PPIs, they are laborious, yield both false positive and false negative interactions, and have so far not generated a comprehensive structural interactome.

To address these challenges, researchers are increasingly using deep learning methods to model protein structures^10^ and PPIs. The most popular predictive algorithm is AlphaFold-multimer (AF-M)^11^, a deep neural network that uses similar principles as AlphaFold to predict structures of multi-chain complexes. These networks achieved state of the art performance by iteratively examining the evolutionary history of proteins from raw Multiple Sequence Alignments (MSAs) along with learned biophysical compatibility to refine and ultimately generate plausible structural predictions. AF-M was trained using five distinct regimens, yielding five models, each of which makes a structure prediction. Many groups are now using AF-M to uncover novel PPIs on the scale of pathways and organisms^12–19^. While many studies have examined AF-M’s ability to correctly model individual protein complexes and predict structures for complexes in curated PPI databases^20–22^, there has been considerably less focus on finding systematic ways to separate true from false interactions in large-scale unbiased screens. In addition, while various groups have proposed different individual metrics for evaluating interface prediction quality and confidence^23^, there has to our knowledge not been a systematic effort to compare them on a single, unbiased dataset or integrate them into a combined and potentially better performing metascore.

We previously used in silico screening with AF-M to uncover how the protein DONSON promotes replication initiation^24^. Folding DONSON with 70 core replication proteins and quantifying the agreement among the five independently trained versions of AF-M (a metric we called average models or “avg_models”) identified five functional DONSON-interacting proteins^25–27^. In the first proteome-wide AF-M screen, we also folded DONSON with over 20,000 human proteins and scored the results using avg_models and other AF-M confidence metrics (ipTM, pDOCKQ)^28^. In this case, DONSON’s functional partners were distributed over the top hundreds or even thousands of hits. Given the poor performance of all existing metrics in this proteome-wide screen, we sought to develop a more robust scoring system that successfully identifies true PPIs.

In this study, we systematically assessed AF-M’s ability to recover true PPIs embedded in a large set of decoy interactions. This analysis showed that standard metrics indeed perform poorly in identifying true interactions. We then used machine learning to train a classifier on a large, curated set of true positive and negative AF-M predictions. This classifier considers structural and biological features of each protein pair and is called SPOC (<u>S</u>tructure <u>P</u>rediction and <u>O</u>mics-based <u>C</u>lassifier). SPOC outperforms standard metrics in separating true positive and negative predictions, including in a proteome-wide in silico screen. We further applied SPOC to an all-by-all interaction matrix of 285 human genome maintenance proteins (40,000 pairs), leading to the identification of many novel, high confidence predictions. These can be viewed and downloaded at predictomes.org, where users can also obtain a SPOC score for their own predictions. In summary, our study provides a well-curated set of AF-M predictions for future machine learning, introduces SPOC, which enables interpretation of large scale in silico screens, and reports a user-friendly web interface that illustrates how large-scale structure prediction in the genome maintenance field drives hypothesis generation.

## Results

### Canonical confidence metrics are inadequate to evaluate large-scale AF-M screens

The results of PPI screens using AF-M are typically ranked using metrics such as the interface <u>p</u>redicted <u>T</u>emplate <u>M</u>odeling score (ipTM, 0-1 scale; Figure S1A), AF-M’s estimate of interface accuracy^29^. Another common metric is pDOCKQ^30^, which considers the predicted number of interacting residues and their local positioning confidence (Figure S1B; 0 - 1 scale). Because pDOCKQ and ipTM scores can be high for structures that contain spurious interfaces (Figure S1C)^14,31^, we previously developed a novel metric that also considers another AF-M output, the predicted alignment error (PAE; 0-30 Å scale, lower is better), a measure of AF-M’s confidence in the *global* positioning of residues. Specifically, we filter AF-M predictions to identify those in which at least one interfacial residue pair involves residues that both have PAE values <15 Å and pLDDT values > 50, and that reside between 1Å and 5Å of each other (Figure S1D). Only protein pairs that satisfy this minimum “contact criterion” (“C+”) are considered. We then quantify what fraction of the C+ residues in a pair are observed in all the independently-trained AF-M models, generating an “average models” (avg_models) score^28^ (Figure S1E; 0-1 scale, > 0.5 is confident). Although this metric was useful in scoring a small-scale in silico screen, its performance was poor in a proteome-wide screen^28^.

To more systematically evaluate the performance of ipTM, pDOCKQ, and avg_models for large in silico screens, we assessed their ability to rank a functional interaction ahead of spurious interactions (“Ranking” experiment; Figure 1A). To this end, we first identified 19 well-characterized protein complexes that were not in the PDB and therefore could not have informed AF-M (Table S1). The only exception was UVSSA/RPB1, whose structure was published after the training cutoff for AF-M v3, which was used for our experiments^32^. For each pair, one protein was selected as “bait” and paired with its correct partner, and it was also paired with 1000 different and randomly selected prey proteins, the vast majority of which should be “true negatives” (TNs; see below). To reduce computation, all pairs were folded in three out of five AF-M models (3 recycles; templates enabled), and each TP and its corresponding 1000 TNs were ranked by various metrics. For example, the bait protein UVSSA was folded with its known partner RPB1 and with 1000 random proteins. The resulting structures were ranked by pDOCKQ, ipTM, and avg_models, which showed that avg_models performed best, giving UVSSA/RPB1 the highest rank (Figure 1B). In a total of 19 ranking experiments, avg_models consistently outperformed pDOCKQ and ipTM. However, in proteome-wide screens, where the number of TNs should be ∼20 times higher than in our ranking experiments, even the avg_models metric would mix the true positive pair with 100-200 random interactions. Indeed, although avg_models performed best in ranking DONSON’s true interactors in a proteome-wide screen^50^, they were still mixed in with hundreds of other proteins, the vast majority of which are presumably spurious hits (Figure S2A-C)^28^. These results show that existing metrics are insufficient to effectively sort the results of large scale in silico interaction screens.

**Figure 1:**
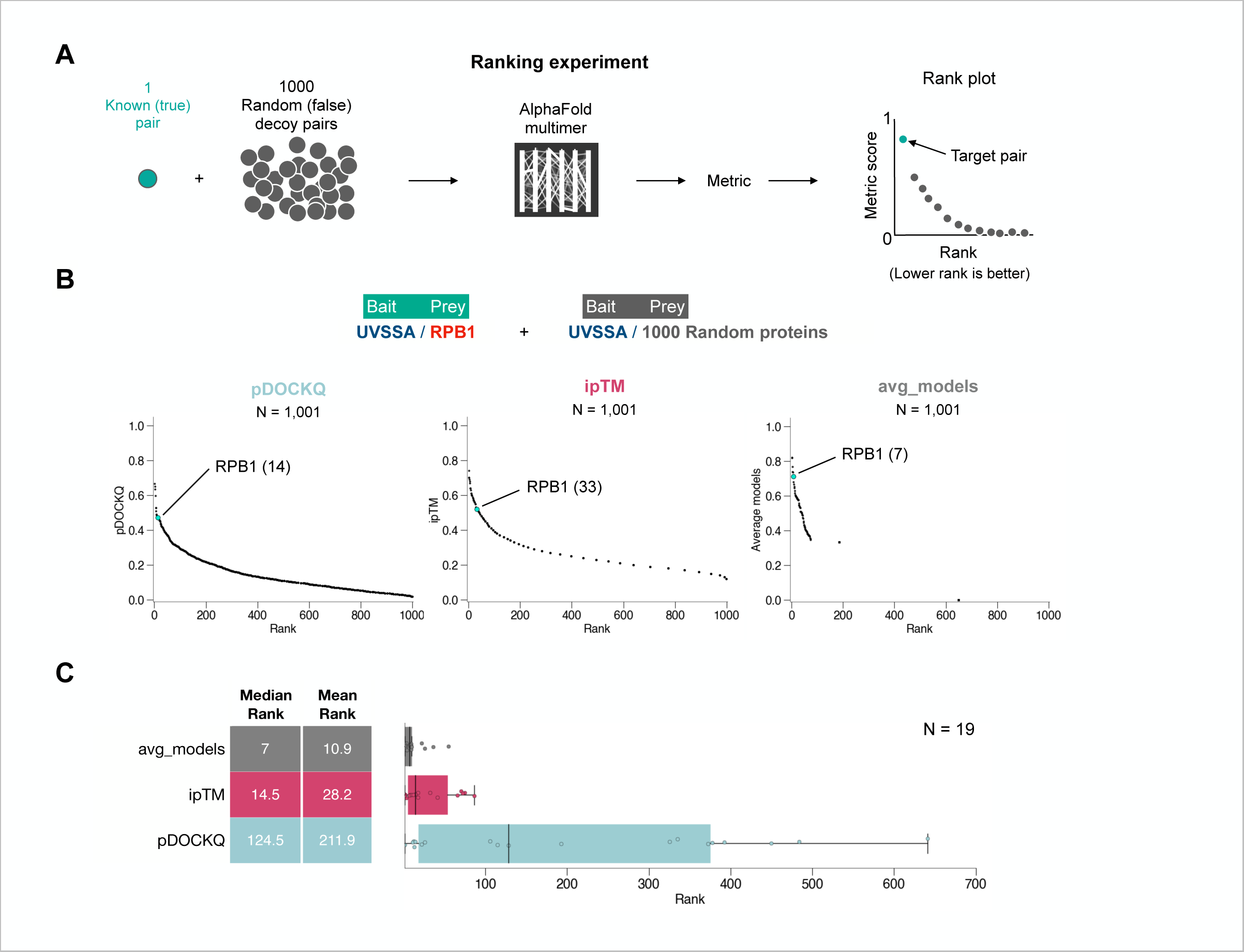
Current metrics are inadequate to evaluate large-scale AF-M screens. **(A)** Schematic of a “ranking experiment” to evaluate AF-M prediction quality metrics. **(B)** Example of a ranking experiment where the target pair UVSSA/RPB1 was embedded in a set of 1000 decoy (UVSSA + random) pairs and evaluated using three metrics. **(C)** Box plots comparing the performance of three different metrics in 19 different ranking experiments. Lines indicate medians and boxes span the 1^st^ to third quartile.

### Curated sets of PPIs to train a new classifier

We sought to use machine learning to train a scoring algorithm or “classifier” that more accurately ranks AF-M predictions. For training and evaluation, we curated four datasets corresponding to biologically meaningful (TPs) and false or spurious interactions (TNs). Careful training set construction was essential because any systematic bias (e.g. differences in protein size between TPs and TNs) would teach the model irrelevant patterns that are not useful in sorting real-world data.

We first constructed the negative set. Previous estimates suggest that the human interactome contains ∼1.5 million TP interactions^33–36^, implying that the vast majority (>99%) of the ∼200 million possible protein pairs correspond to TNs. Therefore, to generate a set in which nearly all pairs are TNs, we folded 40,000 random, binary pairs with our standard folding pipeline. Of the 40,000 pairs folded, 11,993 (30%) satisfied our contact criteria (C+) and were used as the Random Reference Set (RefSet^Random^; Figure 2A). One potential limitation of this approach is that RefSet^Random^ comprises highly dissimilar protein pairs that will not help discriminate against pairs that are in the same pathway or complex but do not interact directly. To address this, we also compiled a negative set of 3,770 “decoy” pairs that reside in well-characterized multi-subunit complexes such as the proteasome or the mitochondrial ribosome but that do not directly interact. 30% (1,115) of these were contact positive C+ pairs and constitute the (RefSet^Decoy^; Figure 2B).

**Figure 2:**
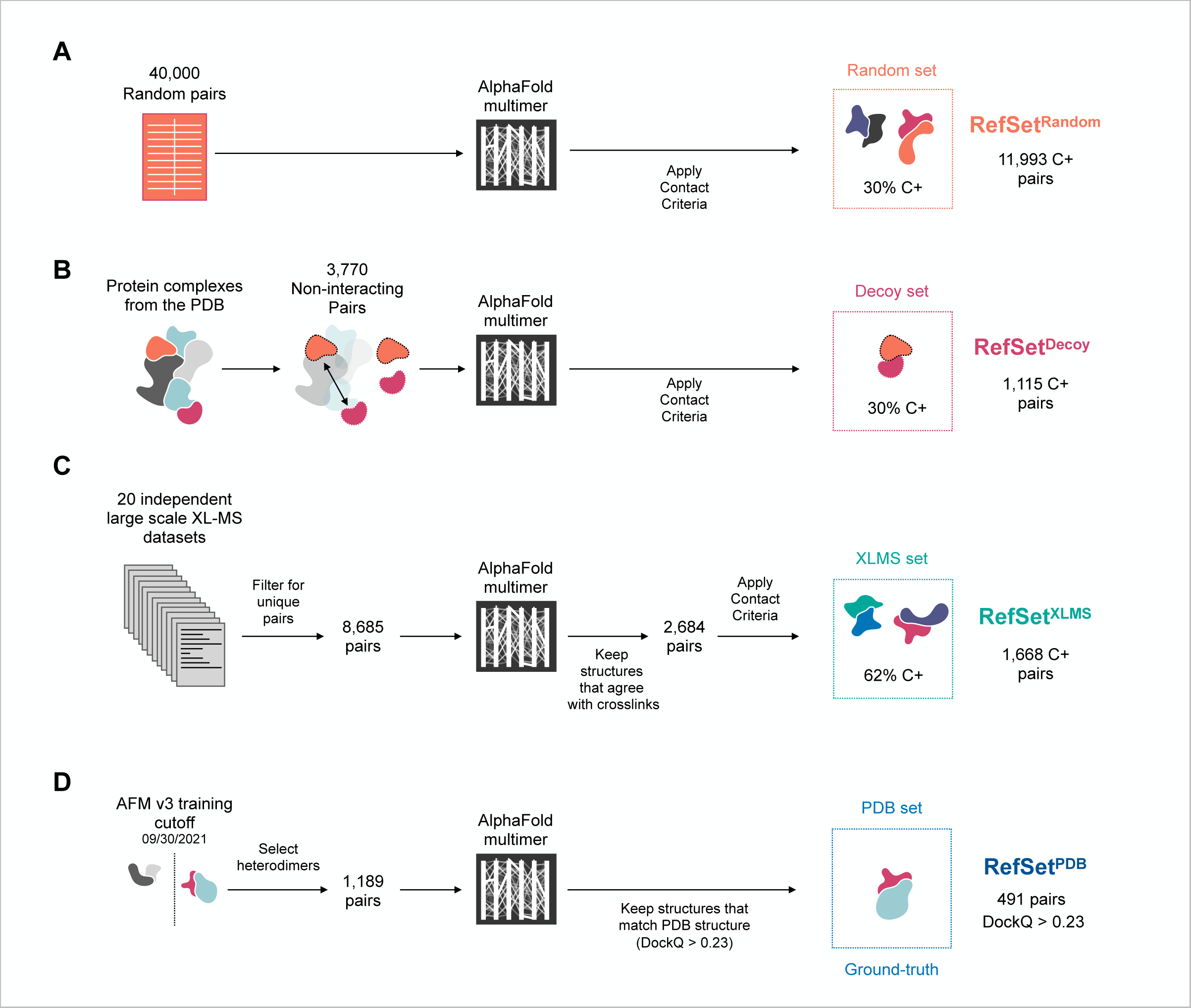
Assembling curated sets for classifier training. **(A)** Schematic illustrating the methodology for constructing the random pairing reference set (RefSet^Random^). We generated 40,000 random pairs by repeatedly sampling from the canonical human protein entries in UniProt. Pairs that exceeded 3,600 amino acids (GPU memory limit) or that were present in another dataset were discarded. **(B)** Schematic illustrating assembly of the PDB decoy set (RefSet^Decoy^). We selected protein pairs that do not make direct contact (defined as more than 10 residue pairs with heavy(non-hydrogen) atoms closer than 5Å) in large multi-subunit complexes of known structure. **(C)** Schematic illustrating assembly of the (RefSet^XLMS^) set, which was mined from human or mouse cross linking datasets. **(D)** Schematic illustrating assembly of the PDB reference set (RefSet^PDB^).

We also sought a true positive reference set that reflects the full spectrum of undiscovered interactions that a screen is intended to uncover. Using only structures from the PDB would likely bias training towards protein complexes that are of current interest and amenable to structure determination. We therefore mined cross-linking mass spectrometry (XLMS) datasets, which capture PPIs in their physiological setting^37^. We compiled cross-linked peptides from 20 XLMS studies (Table S2) and extracted a list of unique binary human pairs. As an important quality control, we only used protein pairs in which at least one of the cross-links detected agreed with at least one of the three AF-M models (residues were predicted to be located within 36 Å, the upper bound of cross-linker lengths used). After folding these 2,684 XLMS pairs, we selected the C+ subset of 1,668 predictions for use as TP pairs (RefSet^XLMS^) (Figure 2C). Importantly, the distribution of protein features (e.g. number of residues, solvent exposed surface area, etc.) for the TP training set (RefSet^XLMS^) and the TN training set (RefSet^Random^ + RefSet^Decoy^) were very similar, except for some differences in protein localization (Figure S3A-B).

Although RefSet^XLMS^ was likely to serve as a diverse TP training set, we could not guarantee that it did not contain false negatives. Therefore, to generate a ground truth dataset for independent evaluation of metrics, we also generated a fourth set of protein pairs corresponding to structures in the PDB that were deposited after the AF-M v3 training cutoff. We then removed any pairs that were already included in previous RefSets. We folded the resulting set of 1,189 heterodimeric pairs with templates disabled to avoid accessing PDB information deposited after training. We then compared our AF-M predictions to the corresponding PDB structures using DOCKQ^38^. Of the 1,064 pairs for which we were able to calculate DOCKQ scores (125 had unresolvable runtime errors), 491 (46.1%) had DOCKQ > 0.23, the CAPRI cutoff for acceptable model quality and are used as a set of ground truth correct pairs (RefSet^PDB^) (Figure 2D).

We used these new, well-curated datasets to further evaluate canonical AF-M metrics. As shown in Figure S2D, 33%, 5%, and 3.4% of the TN pairs (RefSet^Random^ + RefSet^Decoy^) exhibited “positive” pDOCKQ, ipTM, and avg_models scores, respectively, according to standard cutoff values, even though at most 1.5% of these random and decoy pairs should have been positive^33^. In contrast, the RefSet^PDB^ (DOCKQ > 0.23) had many more positive hits, as expected (Figure S2D). However, for all three metrics, there was poor separation between positive and negative pair distributions (Figure S2D). Together, these results are consistent with the poor performance of these metrics in ranking experiments (Figure 1B-C and Figure S2B-C), underscoring the need for a better metric that distinguishes TPs from TNs.

We also asked how existing metrics stratify the XLMS dataset that would be used for training (next section). As expected, pairs from RefSet^XLMS^ displayed higher mean pDOCKQ, ipTM, and avg_models scores than (RefSet^Random^ + RefSet^Decoy^)(Figure S2E). Notably, a substantial number of RefSet^XLMS^ pairs exhibited low pDOCKQ, ipTM, and avg_models scores. It is presently unclear whether this was because RefSet^XLMS^ contains a substantial number of TNs, or because it contains TPs that are particularly difficult to predict and resolve from TNs using conventional metrics.

### Training a classifier that distinguishes TPs from TNs

To develop a classifier that correctly assigns an AF-M structure prediction as true or false, we trained a random forest machine learning model^39^ using the above curated datasets. Random forests use random feature subsets during training to independently build many decision trees that each attempt to assign the correct class to each instance in the training data (Figure S4A). The resulting algorithm is called a classifier. New instances are voted on by all trees, and votes are tallied to produce a classifier score on a 0-1 scale.

To train a “structural” classifier, we developed a series of numeric features that extensively quantify the AF-M predicted interfaces. These included not only AF-M-based metrics (PAE, pLDDT, and avg_models scores), but also many other measurable properties of the interface such as the number of salt bridges and hydrogen bonds between interacting residues (Table S3). After compiling features, we trained the structural classifier by randomly splitting our TN (RefSet^Random^ + RefSet^Decoy^) and TP (RefSet^XLMS^) datasets into training (75%) and testing groups (25%). We did not train on RefSet^PDB^ to preserve this dataset for subsequent classifier analysis (see below), and to avoid bias. After fitting a random forest using the training data and features, we evaluated its performance, as well as the performance of previously developed metrics, on the 25% testing data held back. Performance was assessed in Recall and <u>F</u>alse <u>D</u>iscovery <u>R</u>ate (FDR) curves (Figure 3A). These curves quantify how many TPs and TNs are identified above each classifier threshold and display the results as the Recall rate (Figure 3A, solid lines; fraction of all TPs that are captured above the threshold) and FDR (Figure 3A, dotted lines; fraction of all pairs captured above the threshold that are TN). As a single measure of performance, we determined the Recall rate when the FDR was 5% (1 in 20 interactions labeled by the classifier as true are actually false). Compared to existing metrics, the structural classifier exhibited the highest recall (0.76) at 5% FDR (Figure 3A; see table). We also evaluated performance using Receiver Operating Characteristic (ROC) curves (Figures S4B), which quantify the TP and FP rates as a function of classifier score. Using this metric, the structural classifier also performed best, achieving the highest <u>A</u>rea <u>U</u>nder the ROC <u>C</u>urve (AUC) of 0.90.

**Figure 3:**
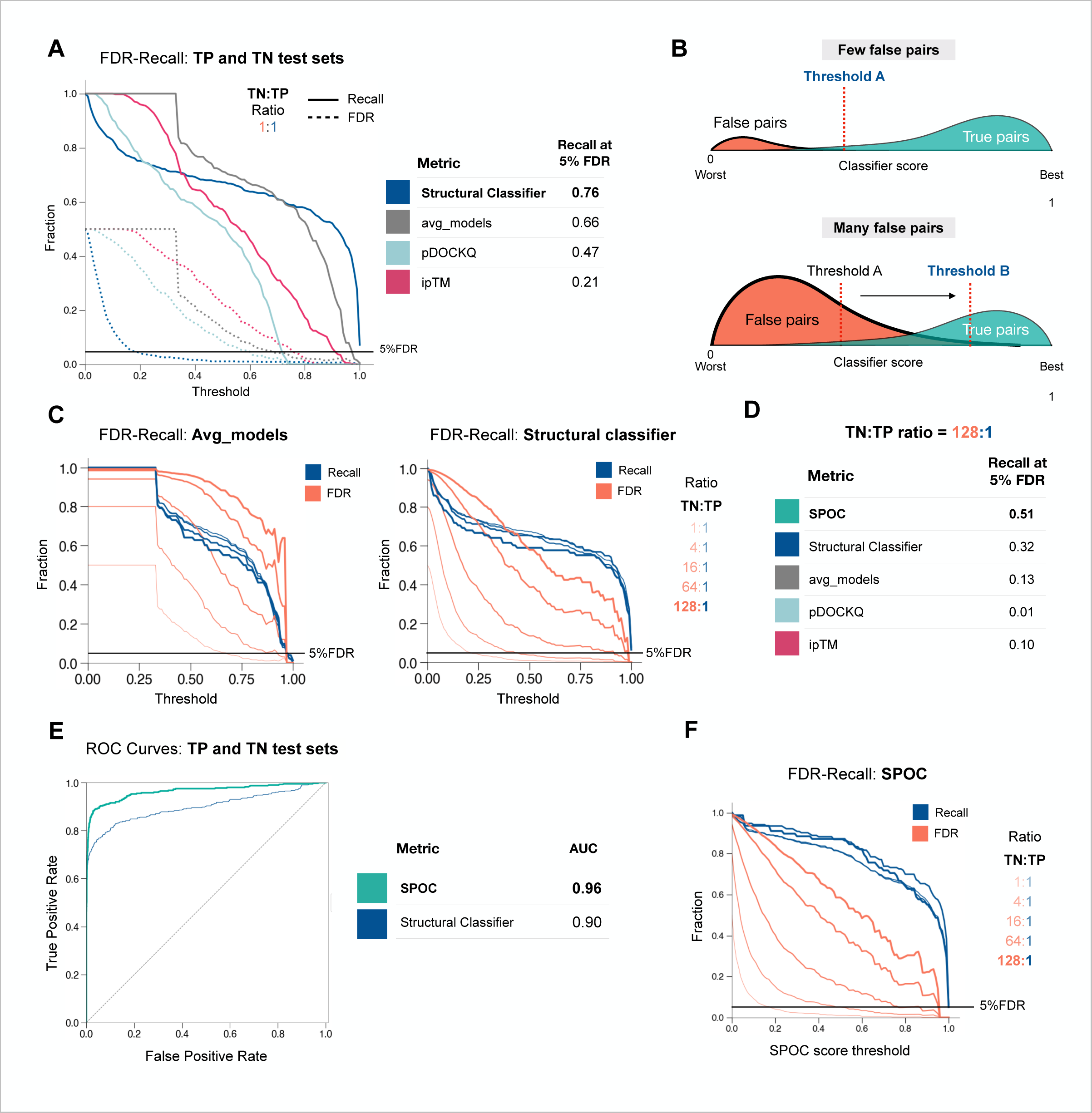
Classifier performance on curated test sets. **(A)** A plot of Recall (solid lines) and FDR (dotted lines; same as 1 - precision) as a function of selected threshold for three previously used metrics and the new structural classifier at a TN:TP ratio of 1:1. TN is (RefSet^Random^ + RefSet^Decoy^); TP set is RefSet^XLMS^. **(B)** Schematic illustrating that as the abundance of false pairs rises, a higher classifier threshold is required to maintain a low FDR. **(C)** Recall-FDR plots for avg_models and the structural classifier at various TN:TP ratios. TN is (RefSet^Random^ + RefSet^Decoy^); TP set is RefSet^XLMS^. **(D)** Table comparing the recall fraction at 5% FDR and TN:TP = 128:1 for avg_models (panel C), structural classifier (panel C), and other metrics (curves not shown). **(E)** AUC under the Receiver Operating Characteristic (ROC) curves for SPOC and the structural classifier. TN is (RefSet^Random^ + RefSet^Decoy^); TP set is RefSet^XLMS^. **(F)** Recall-FDR plots for SPOC at various TN:TP ratios. TN is (RefSet^Random^ + RefSet^Decoy^); TP set is RefSet^XLMS^.

Like many other studies, the above evaluation of classifier performance involved test sets with equal numbers of TNs and TPs (reviewed in ^40^). However, this 1:1 TN:TP ratio does not accurately reflect real-world scenarios, the most extreme of which occurs when a single protein is screened against the entire proteome. In this case, we estimated the TN:TP ratio to be ∼80:1 (Figure S3C). In general, as the ratio of TNs:TPs increases, so does the number of TN pairs with high scores (false positives); due to this “invasion” by TNs, the threshold must be increased to maintain an acceptably low FDR (Figure 3B). Therefore, to obtain a more realistic estimate of classifier performance, we created test sets spanning TN:TP ratios from 1:1 to 128:1. As expected, the Recall rate was largely unaffected by these ratios, but the FDR increased progressively as the proportion of TNs increased (Figure 3C). At the 128:1 ratio and 5% FDR, the structural classifier retained the highest recall (32%) as compared to avg_models, pDOCKQ, and ipTM (Figure 3C-D). Analyzing classical precision recall curves similarly demonstrated that the structural classifier maintained the highest recall and precision at high TN:TP ratios (Figure S4C). Therefore, the structural classifier outperformed other metrics under proteome-wide screening conditions.

While the structural classifier performed favorably relative to other metrics, its 32% recall at high TN:TP ratios was unsatisfactory. We therefore attempted to train a better classifier by including genome-wide “biological” features of each protein being queried that were external to AlphaFold, including data from DEPMAP^41^, coexpressionDB^42^, PerturbSeq^43^, T5 protein language model embeddings^44^, subcellular colocalization predictions from DeepLoc 2.0^45^, hit profiles from the BioORCS CRISPR screen database, and high throughput interaction experiments from the BioGrid database^46^ (Table S3). Given its biased nature, we did not include information from the protein association database STRING, or other researcher-curated resources. We then re-trained and evaluated a new <u>S</u>tructure <u>P</u>rediction and <u>O</u>mics informed <u>C</u>lassifier (SPOC) and found that it significantly outperformed all other algorithms, including the structural classifier: its AUC in ROC curves was 0.96 (Figure 3E), and at the 128:1 TN:TP ratio, it recalled 51% of positive pairs at 5% FDR (Figure 3D and 3F). This is nearly two times higher recall than the structural classifier, and at least ∼4 times better than all other metrics. Strikingly, at a threshold of 0.97, SPOC achieved an effective FDR of 0% while still recalling ∼50% of TPs, even at the highest TN:TP ratio (Figure 3F).

Many different features contributed to SPOC’s output, with no single feature dominating (Figure S4D). As expected from the fact that SPOC performed better than the structural classifier, both “biological” and “structural” features were assigned high importance scores (Figure S4D). Our results suggest that in a curated dataset, SPOC identifies true interactions with impressive sensitivity and high specificity, even at TN:TP ratios that approximate proteome-wide screens.

### Evaluating classifier performance in biological discovery

To examine SPOC’s performance using an orthogonal performance test, we revisited its ability to rank functional interactions ahead of spurious ones (Figure 1A). Consistent with its ability to analyze curated datasets (Figure 3E), SPOC outperformed all other metrics in these ranking experiments (median rank = 1, mean rank = 1.9)(Figure 4A) (Table S1). Importantly, it performed well on proteins in different compartments and pathways, as expected given that training was pathway- and compartment-agnostic (Figure S5C and Table S1). Moreover, whereas conventional metrics generally distributed pairs evenly across their respective score ranges, SPOC more clearly separated the TP from the TNs (Figure 4B, Figure S5A-C; Note that the classifier’s top hit in the SLF1 mini-screen was RAD18, a SLF1 interactor^47–49^. Finally, we assessed how SPOC ranked DONSON’s five interactors (MCM3, SLD5, TOPB1, DPOE2, and DONSON; Figure S2A) in our previous proteome-wide screen for DONSON interactors^50^. Unlike all other metrics (Figure S2B-C) including the structural classifier (Figure 4C), SPOC placed DONSON’s functional partners in the top 7 hits out of more than 20,000 pairs (Figure 4D). This was remarkable given that DONSON was not present in any of the training data. These results show that SPOC outperforms all other metrics in real-world ranking experiments, and can discover PPIs *ab initio* in proteome-wide in silico screens.

**Figure 4:**
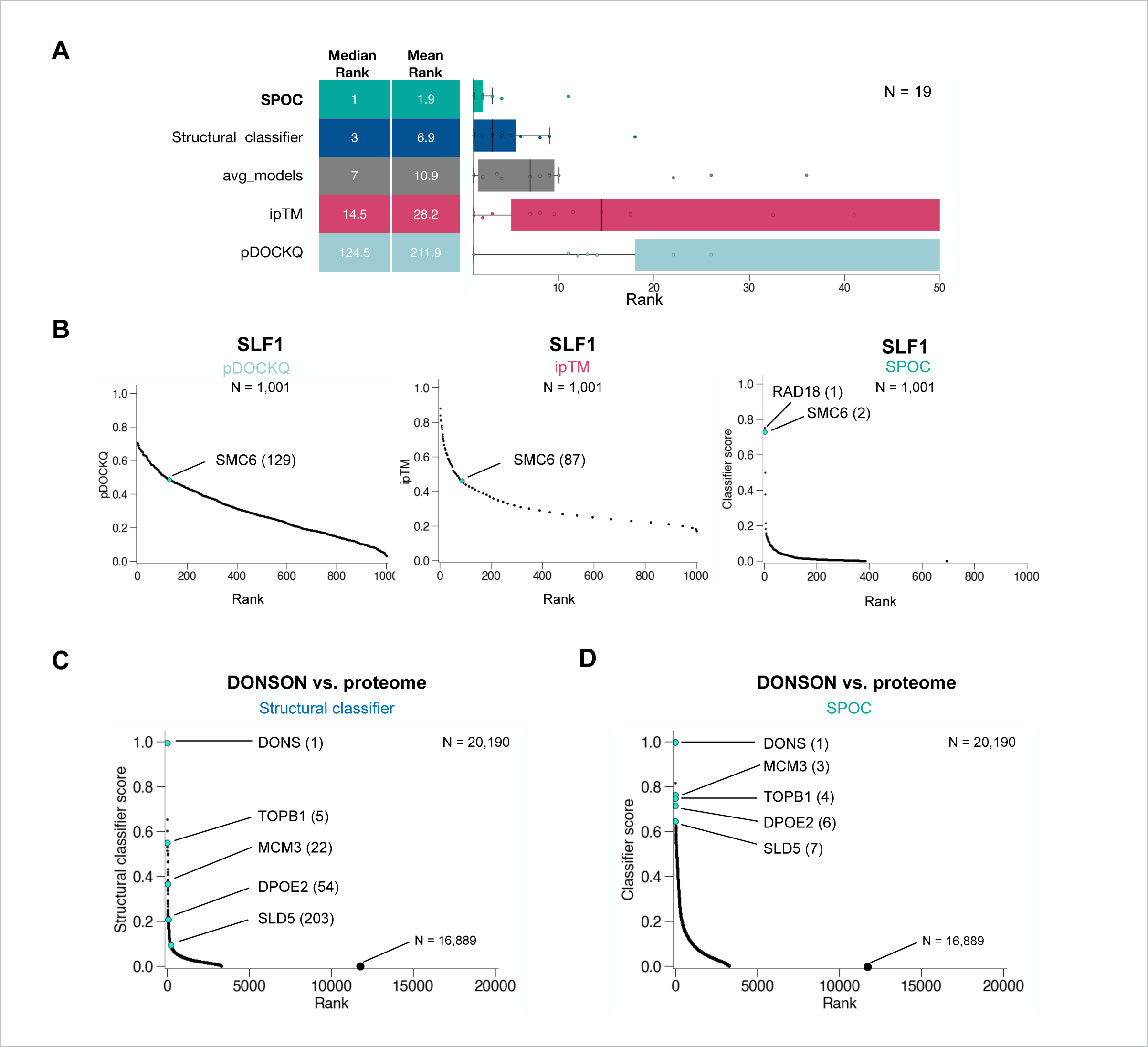
Evaluating SPOC performance for biological discovery applications. **(A)** Box plots comparing the performance of five ranking metrics. The data for avg_models, ipTM, and pDOCKQ are the same as in Figure 1C. X-axis was truncated above rank 50. **(B)** The SLF1/SMC6 true pair was embedded in 1000 random pairs involving SLF1, and the AF-M predictions for each pair were ranked using three different metrics. **(C)** DONSON was folded with more than 20,000 human proteins, and the resulting predictions were ranked using the structural classifier. True DONSON interactors are indicated in cyan. **(D)** Same as (C) but using SPOC for ranking.

### In silico screening in genome maintenance

Having developed SPOC, we sought to comprehensively predict all possible pairwise interactions within a biological pathway. We therefore compiled a list of 285 core human genome maintenance (GM) proteins and folded all binary combinations of these factors in three out of the five AF-M models, yielding structure predictions for more than 40,000 protein pairs. Of the 40,174 GM pairs, 13,175 (32.8%) were C+, satisfying our contact criteria. To find a suitable SPOC score cutoff for positive interactions in this dataset, we plotted the F1 score (optimal balance between recall and precision) as a function of the SPOC score, which peaked at a SPOC value of 0.3 (Figure 5A). On our curated positive test set (RefSet^PDB^), this score cutoff captured 88% of TPs and 2.1% of TNs from the (RefSet^Random^ +RefSet^Decoy^) (Figure 5B). Across all genome maintenance pairs, 2,093 (5.2%) had SPOC scores > 0.3 (Figure 5C), implying that each GM protein might have on average seven true partners in the matrix.

**Figure 5:**
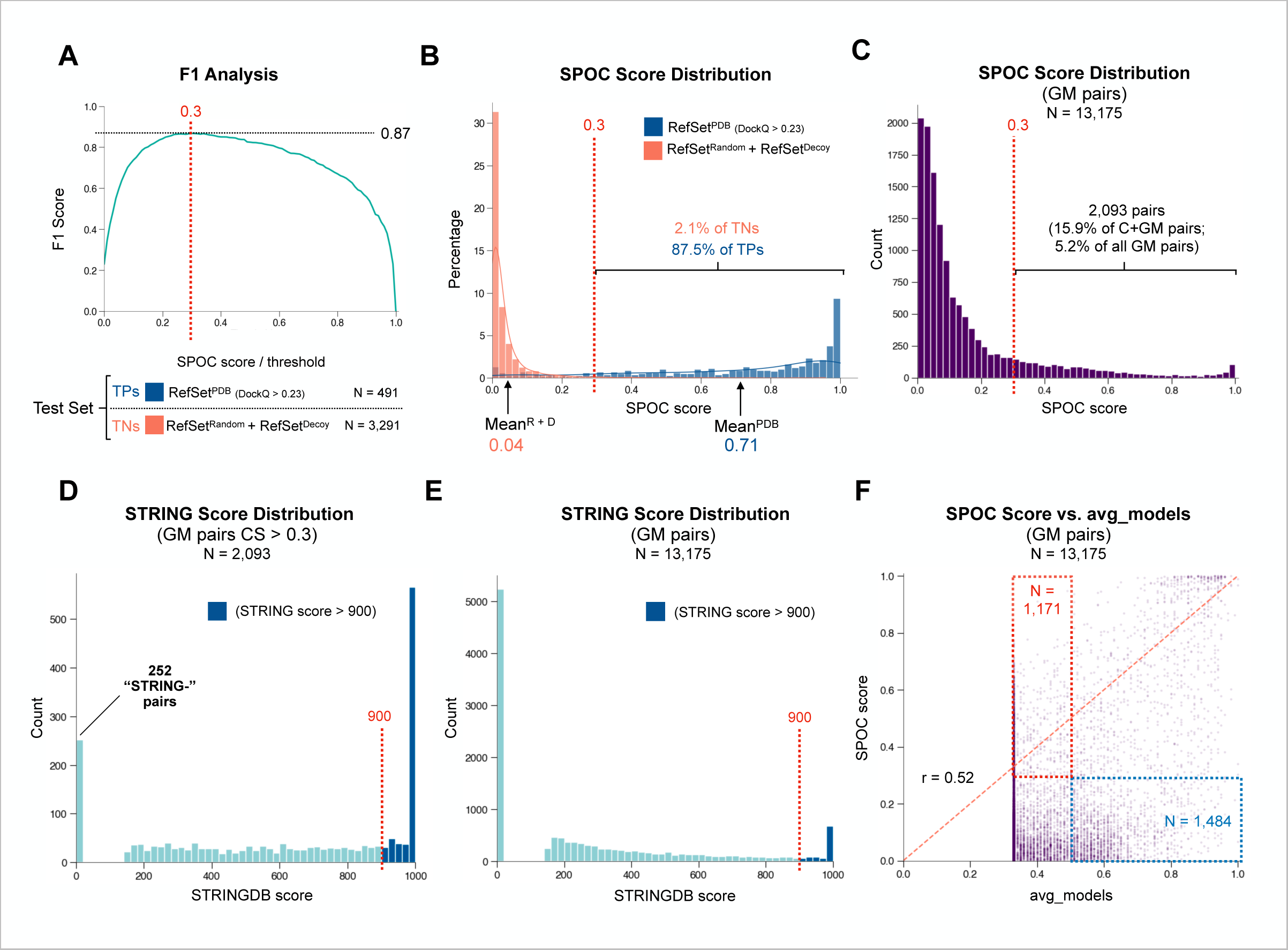
Applying SPOC to find novel interactions in a biological pathway. **(A)** A plot of the F1 (TP/(TP + 0.5*(FP + FN)) score as a function of SPOC score/threshold applied to the TN (RefSet^Random^ + RefSet^Decoy^) and TP (RefSet^PDB^; DockQ > 0.23) test sets. The F1 score peaks at a threshold SPOC score of 0.3. **(B)** Histogram showing the distribution of SPOC scores for TNs (RefSet^Random^ + RefSet^Decoy^) and TPs (RefSet^PDB^; DockQ > 0.23). A proposed cutoff for true interactions (0.3), as well as the % of TPs and TNs above the cutoff are shown. **(C)** The distribution of SPOC scores on 13,175 C+ <u>G</u>enome <u>M</u>aintenance (GM) pairs. 15.9% of these interactions achieved a SPOC score > 0.3. **(D)** Histogram of STRING DB scores associated with the 2,093 pairs with SPOC score > 0.3 shows that 723 (34.5%) top classifier scoring interactions also have high (>900) STRING scores. **(E)** Histogram of STRING DB scores for all 13,175 C+ GM pairs. 5,240 pairs have scores of 0, indicating that no prior text or data has suggested a potential association while 978 (7.4%) have scores greater than 900. **(F)** A scatter plot comparing the SPOC score (y) to the avg_models score (x) for the 13,175 C+ pairs in the GM dataset. The dashed orange line shows how a perfect correlation between the two scoring schemes would look. See text for explanation of red and blue boxes.

We addressed how SPOC compares to curation by STRING, in which association strength scores exceeding 900 (on a 0-1000 scale) indicate a strong physical or biological interaction. Notably, 723 of the 2,093 GM pairs (34.5%) with a SPOC score > 0.3 had STRING scores >900 (Figure 5D), compared to 978 among all 13,175 pairs (7.4%) in the GM group (Figure 5E), a ∼5-fold enrichment. Interestingly, 252 of the 2,093 pairs with high SPOC scores (12%) were absent from STRING, representing potential PPIs for which there was no documented association (Figure 5D; Table S4). The top pair is MMS22L-RPA2, which is consistent with prior evidence that the MMS22L-TONSL complex binds to RPA-ssDNA filaments^51^. The second highest pair is USP37-CDC45, which is consistent with our previous finding that USP37 interacts with CMG complexes^52^. We also addressed how SPOC and the best prior metric, avg_models, compare in the analysis of the GM data. Plotting the SPOC score vs. avg_models showed some correlation (r = 0.52), but overall low agreement (Figure 5F). Importantly, there were many pairs with low avg_models scores (<0.5) that SPOC rated highly (>0.3; red box) and conversely, many pairs with high avg_models scores (>0.5) that SPOC downgraded (<0.3; blue box). These results underscore that SPOC makes substantial adjustments to previous rankings.

The GM group contains many PPIs with high confidence scores that are not structurally resolved but are nevertheless supported by strong biochemical or genetic evidence, indicative of SPOC’s ability to detect meaningful interactions. In many cases, the PPI has been mapped sufficiently to indicate that the structure prediction is correct. An example is the CIP2A-TOPBP1 pair (SPOC=0.799), in which residues experimentally shown to be critical for the interaction^53^ agree with the AF-M prediction (see predictomes.org). Other examples include CIP2A-CIP2A (SPOC=0.993; ^54^), FIGNL1-FIRRM (SPOC=0.995;^55,56^), and MMS22L-TONSL (SPOC=0.985;^57–59^)(Table S4). This “retroactive validation” suggests that the GM dataset contains many valid predictions, and that in silico screening is a powerful approach to detect new PPIs.

### A web portal for AlphaFold multimer predictions

To allow researchers to interact with the genome maintenance structure prediction data, we created predictomes.org, a user-friendly online database (first released in September 2023). Users can browse an expandable, interactive matrix (Figure 6A) or a sortable list (Figure S6A) that can be ranked by SPOC score and other metrics. Clicking on a matrix tile or a list entry displays an information page that includes an interactive protein structure viewer (Figure 6B)^60^ from where the AF-M structure predictions can be downloaded. If experimental structures of the pair already exist, the corresponding PDB entries are listed above the structure viewer (Figure 6B; blue arrow), and the PDB structures can be superimposed on the AF-M prediction (Figure 6B; red arrow). The information page also contains UniProt entry information, residue level evolutionary conservation, predicted residue contacts, interactive PAE and pLDDT plots, and data from the STRING and BioGRID databases about potential associations (Figure S6B and predictomes.org). These features allow rapid visualization, ranking, and triage of thousands of structure predictions.

**Figure 6:**
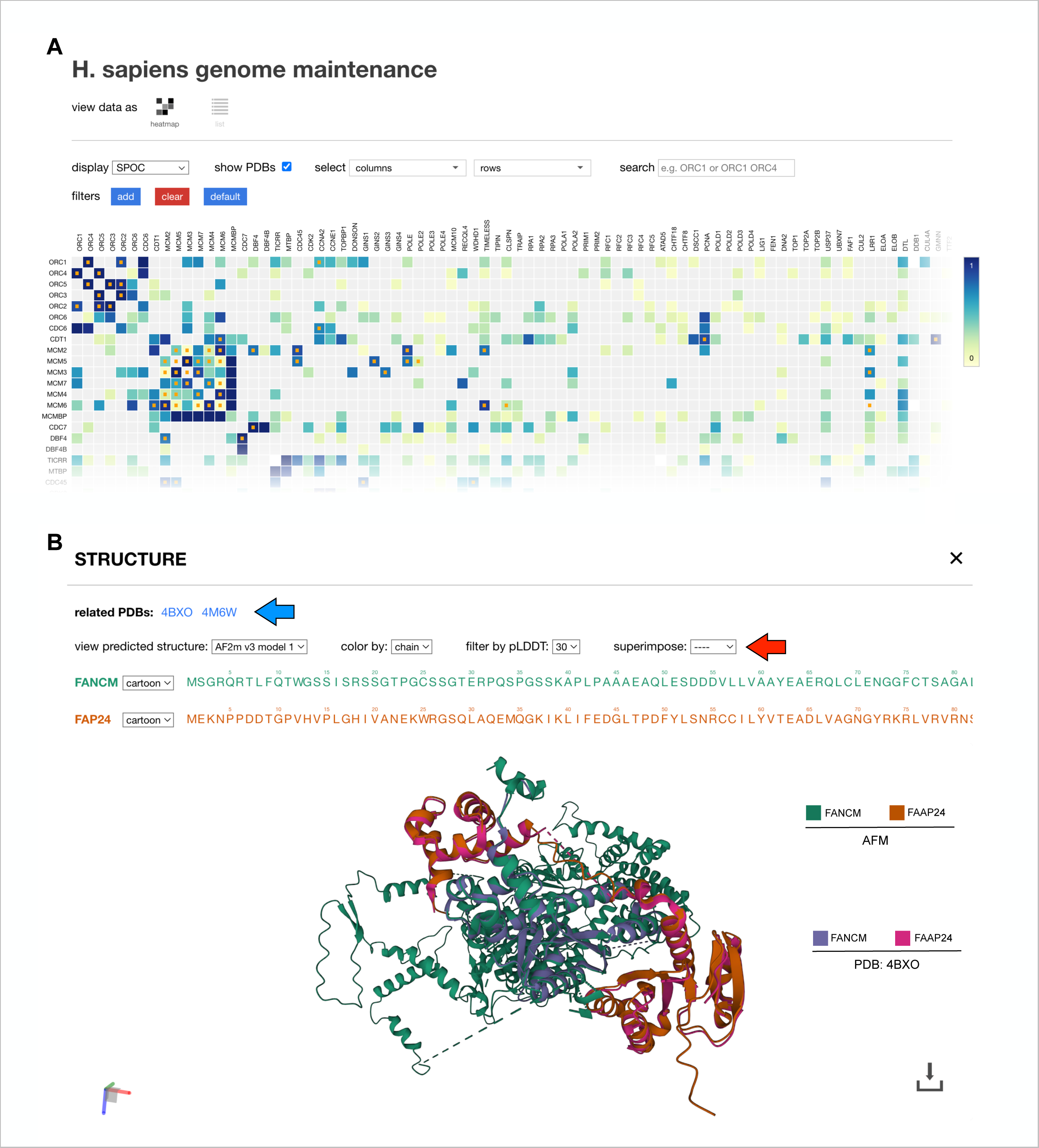
A web portal for AF-M predictions. **(A)** Screenshot of the interactive matrix from predictomes.org. Tile color darkness is proportional to the displayed confidence metric (SPOC). Tiles with orange dots represent pairs that are found in the PDB. Specific biological pathways can be selected for display in rows and columns. **(B)** Screen shot of the interactive structure viewer for the FAAP24-FANCM pair. The FANCM-FAAP24 structure (PDB 4BXO; purple and pink chains) was superimposed on the AF-M structure prediction (green and orange chains) using the superimpose tool (red arrow). There are different options to display the structure, filter residues by pLDDT, and color the structures by different metrics such as pLDDT.

### AI-driven hypotheses generation

The genome maintenance predictome contains many new, high confidence predictions that suggest interesting and testable hypotheses. We highlight two examples related to replicative DNA polymerases. The first involves lagging strand synthesis (Figure 7A). In this process, DNA polymerase α (pol α), which interacts stably with the CMG replicative helicase ^61^, first primes each new Okazaki fragment^61^ on the lagging strand template. RFC then loads the processivity factor PCNA on these primers, followed by primer extension by pol δ. Interestingly, AF-M predicted with high confidence (SPOC=0.921) that the non-catalytic POLD3 subunit of DNA pol δ extends an exposed beta sheet in the catalytic POLA1 subunit of pol α (Figure 7B and S7A; Table S5 for other confidence metrics). This interaction was predicted from humans to fission and budding yeasts (Table S5), and in budding yeast, previous experiments mapped this interaction to the location in both proteins predicted by AF-M^62,63^. Interestingly, the same region of POLA1 was also predicted to bind a small peptide in the N-terminal unstructured region of RFC1 (SPOC=0.789), the largest subunit of the RFC complex (Figure 7B and Figure S7B; Table S5). Strikingly, in three-way structure predictions, the POLD3 and RFC1 peptides interacted with each other on the surface of POLA1, with RFC1 draping over the composite beta sheet formed by POLA1 and POLD3 (Figure 7B). This ternary complex was predicted with high confidence across metazoans and fission yeast but not in budding yeast (Table S5; SPOC only scores binary predictions), and it is consistent with isolation of a pol δ - pol α -RFC complex from mammalian cells^64^. These predictions suggest that when pol α primes a new Okazaki fragment, RFC and pol δ are already attached to pol α, allowing seamless transfer of the primer from pol α to RFC to load PCNA, followed by engagement of pol δ and primer extension (Figure 7A). In agreement with this idea, single molecule experiments demonstrate that yeast pol δ remains bound to the replisome over multiple cycles of Okazaki fragment synthesis, an effect that depends on Pol32, the yeast counterpart of POLD3^65^.

**Figure 7:**
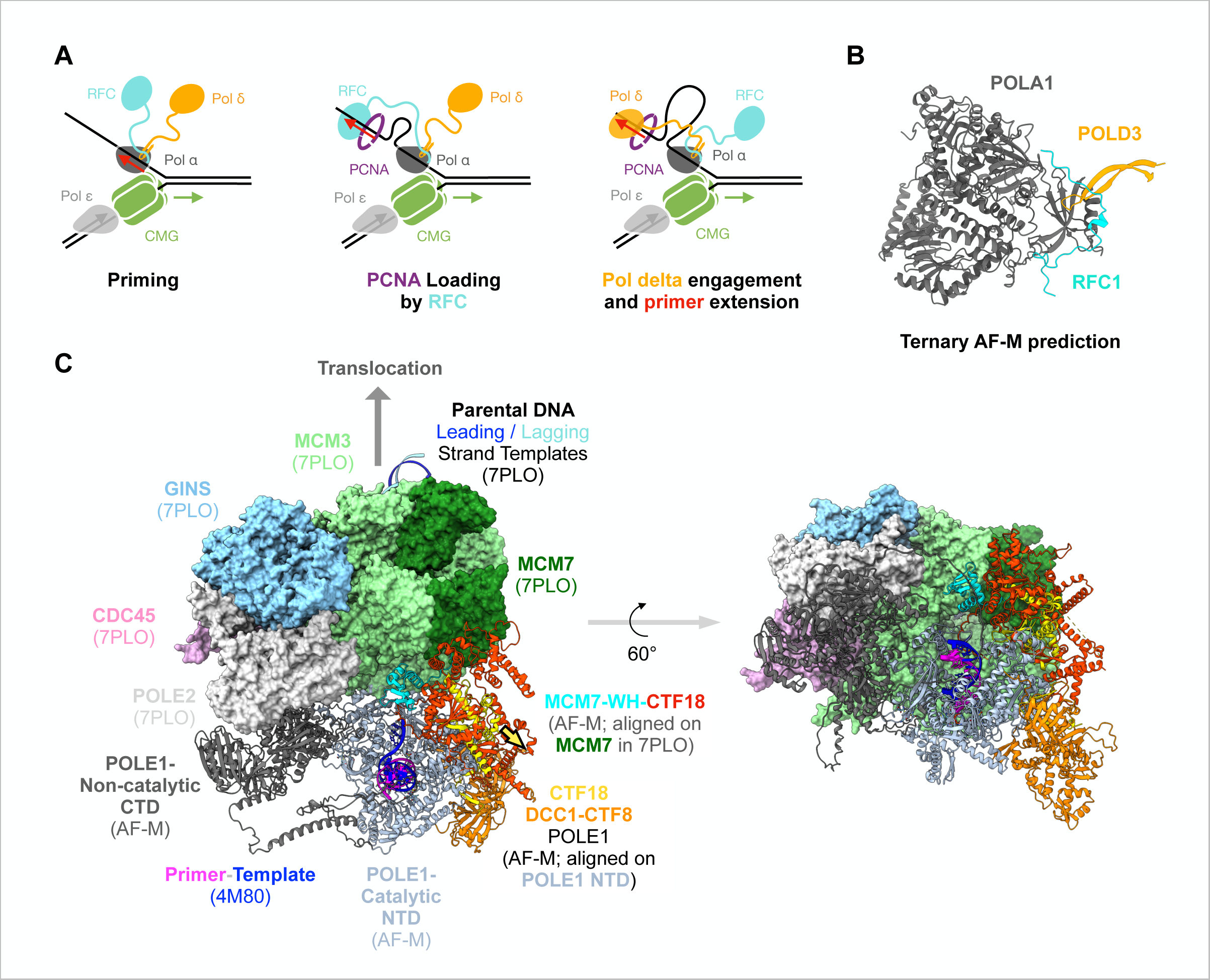
New hypotheses suggested by predictomes.org. **(A)** Model of processive Okazaki fragment processing. During Okazaki fragment priming by pol a, pol d (via POLD3) and RFC (via RFC1) bind cooperatively to pol a (via POLA1). As soon as pol a releases the primer, the tethered RFC occupies the primer-template and deposits PCNA. When RFC dissociates, the tethered pol d occupies the primer-template and initiates primer extension. **(B)** AF-M prediction of POLA1 (grey; residues 310-1264), POLD3 (yellow; residues 380-412), and RFC1 (cyan; residues 153-190) folded at the same time. For POLA1-POLD3 and POLA1-RFC1 binary predictions, see figures S7A-B and predictomes.org. **(C)** AF-M-informed model of how the POLE1 catalytic domain is positioned near CMG’s exit channel by the interaction of CTF18 (red) with the winged-helix domain of MCM7 (cyan) and the POLE1 catalytic domain. The model was assembled as follows: An AF-M prediction of POLE1 (C-terminal non-catalytic domain shown as grey ribbon, N-terminal catalytic domain shown as light blue ribbon) was aligned on the C-terminal, non-catalytic lobe of POLE1 in the cryo-EM structure of the human replisome (PDB: 7PLO, of which only MCM2-7, CDC45, and GINS are shown, POLE1 hidden). To model the primer template, the structure of yeast POLE1 catalytic domain with a primer template (PDB: 4M8O) was aligned on the POLE1-NTD shown (yeast POLE1 hidden; primer-template shown). We also generated an AF-M prediction of a complex of CTF18, CTF8, DCC1, and the NTD of POLE1 (which matches key features of an analogous experimental structure, PDB 6S2E;^73^), and aligned it on the POLE1-NTD shown. This revealed that the CTF18 (yellow)-DCC1(orange)-CTF8(orange) complex binds the distal side of POLE1-NTD. Separately, MCM3, MCM7, and CTF18 were folded together and aligned on MCM7 from 7PLO. This shows that a movement of 30Å (yellow arrow) would superimpose the CTF18 aligned on MCM7 (red) and the CTF18 aligned on POLE1-NTD (yellow). Given the reported flexibility between the NTD and CTD of POLE1^74^, and some predicted flexibility between the NTD and CTD of CTF18 (https://alphafold.ebi.ac.uk/entry/Q8WVB6), CTF18 should be able to bind MCM7 and POLE1-NTD simultaneously. In this way, CTF18 would tether the POLE1-NTD near the rear exit channel of CMG, with the leading strand template (dark blue strand) being fed into the active site.

The second novel hypothesis addresses how the leading strand DNA polymerase ε (pol ε) and the CTF18 complex are oriented on the replisome. In particular, the location of the catalytic domain of pol ε on the replisome remains mysterious. AF-M predicted an interaction between CTF18 and the winged helix domain of MCM7 (SPOC = 0.579), which resides near the rear exit channel of CMG (Figure 7C, red and cyan ribbon diagrams; Table S5). Together with the extensive interface between the CTF18 complex and POLE1^66^, these interactions would position the catalytic, N-terminal domain of pol ε adjacent to CMG’s rear exit channel (Figure 7C, grey ribbon diagram; see figure legend). In this way, the leading strand template (Figure 7C, blue strand) would be fed directly into the pol ε active site.

The above examples illustrate how large-scale screening for binary PPIs uncovers new predictions that seed novel mechanistic hypotheses, some of which are readily aligned with existing data.

### SPOC Tool

We wanted to facilitate broad access to SPOC. We therefore trained a computationally less intensive classifier model (compact SPOC, “cSPOC”) in which we reduced the number of decision trees from 1,000 to 500 and removed features that did not significantly contribute to classifier output. cSPOC demonstrated nearly identical performance as the original when measured by AUC in ROC curves (Figure S6C) and recall-FDR rates (Figure S6D). We created an online tool at predictomes.org that uses cSPOC to analyze up to 50 user-generated AF-M structures at a time. After uploading their predictions, users are sent cSPOC, ipTM, pDOCKQ, and avg_models scores for each protein pair. We expect that cSPOC access will broadly facilitate identification of biologically plausible structure predictions.

## Discussion

Here, we report tools and resources that will help biologists leverage the structure prediction revolution for their research. First, we generated well-curated sets of true positive and true negative protein pairs that can direct future machine learning efforts. Second, we used these datasets to train SPOC, a new classifier that effectively discriminates biologically meaningful and spurious AF-M predictions, even under the most demanding scenario of a proteome-wide screen. We make a simplified version of this classifier (cSPOC) available as an online tool to allow classification of user-generated structure-predictions. Third, we generate a SPOC-curated structural predictome of genome maintenance proteins, and we give examples of how it can be leveraged to seed new hypotheses. Together, the work helps lay the foundation for the eventual development of a comprehensive structural interactome.

Two independent forms of evidence show that SPOC outperforms all previous metrics. First, on curated testing sets containing TPs and TNs, SPOC exhibits the highest AUC values in ROC curves (0.96 vs. next best of 0.9), and it captures the largest number of true positive interactions under realistic screening scenarios where TNs greatly outnumber TPs (51% vs. the best conventional metric of 13%). Second, in orthogonal performance tests involving real in silico screens, including one that was proteome-wide, SPOC ranked true pairs higher than any other metric.

An important consideration is how to use SPOC to identify positive pairs while minimizing false positives. As shown in Figure 5B, positive pairs can in principle have any SPOC score, but they are strongly enriched for high scores. Therefore, the appropriate SPOC threshold depends on context and what FDR is tolerable, which can be judged from the Recall-FDR curves for SPOC (Figure 3F). A proteome-wide screen is expected to involve an ∼80:1 TN:TP ratio and is therefore best modeled by the 64:1 and 128:1 curves, where maintaining a 5% FDR requires a classifier threshold of ∼0.95. In contrast, looking for direct interactors in a mass spectrometry pulldown is more appropriately evaluated at a lower ratio such as 16:1 or 4:1, where 5% FDR is compatible with thresholds of ∼0.5-0.75. Invariably, the safest strategy is to analyze interactions from highest to lowest SPOC score. Although a SPOC score of >0.97 yielded an apparent FDR=0 in curated datasets, even at the 128:1 ratio (Figure 3F), it is uncertain whether this holds true for real-world data. In other words, a high score alone never provides proof of an interaction, and conversely, a low classifier score is not proof that an interaction is false. We believe that when used judiciously, SPOC provides a powerful tool to prioritize hits and drive mechanistic discovery.

It is interesting to consider the source of erroneous classifications in AF-M screens. False positives might arise because AF-M was only trained on true positives from the PDB and therefore attempts to find a solution for all pairs. Another possibility is that some false positives would in fact interact if brought together in vitro (“biophysical interactors”). Physical interaction studies on such pairs will be required to distinguish these possibilities. Many false negatives probably involve protein pairs that are scaffolded by other factors and whose interaction is too minimal to be detected by AF-M in binary screens. Other false negatives probably arose from our AF-M protocol, which was chosen to maximize throughput. Many of these can probably be eliminated by increased sampling^67^ or segmenting proteins^31,68^.

SPOC likely has “blind spots.” These will include PPIs with physical features that are not well represented in the XLMS data on which the classifier was trained, as well as pairs that do not exhibit biological patterns (e.g. co-expression, co-localization, genetic co-dependence) typically associated with interacting proteins. An example of the latter instance would be a PPI in which the two interacting proteins’ major functions and partners are in orthogonal pathways. One way to better detect the latter groups is to identify more informative structural features of PPIs to improve classifier training. Another is to further improve structure prediction algorithms themselves so that their predictions can be trusted without external biological data.

A more compact version of SPOC (cSPOC) is accessible at predictomes.org and will calculate scores for researcher-generated AF-M predictions. This tool works best when applied to predictions generated using AF-M settings that resemble as closely as possible those used to train the classifier. Accordingly, if a user uploads more than 3 predictions per pair, the tool randomly analyzes three to mirror our training regimen. However, other AF-M settings such as the number of recycles or dropout enabling that cannot be adjusted post-run may impact predictions. In many cases, it will therefore be advisable to regenerate and analyze AF-M structures using the same protocol used for classifier training.

The data in predictomes.org catalyzes the formulation of novel hypotheses. By sorting each protein’s putative partners as a ranked list and displaying predictions in an interactive structure viewer with relevant, accompanying information, users can rapidly triage vast numbers of structure predictions and thereby identify the most promising interactions. Together with second generation structure predictions, this resource allows the formulation of compelling new hypotheses. An example is our finding that POLA1 is predicted to interact with POLD3 and RFC1. This observation motivated a secondary folding experiment involving all three proteins, which predicted the existence of a ternary complex of pol δ, RFC, and pol α that promotes Okazaki fragment in a processive assembly line, consistent with single molecule experiments^65^. We believe that generating, organizing, and curating structure predictions in major biological pathways, and eventually proteome-wide, will launch a new era of accelerated mechanistic discovery in the biological science.

## Acknowledgements

We thank Alan Brown, Lucas Farnung, Wade Harper, Edward Huttlin, Nicholas Polizzi, and members of the Walter laboratory for helpful discussions and comments on the manuscript. We thank Jack Shaw, Helen Zhu, and Maksym Shyian for support in writing helpful analysis scripts. E.W.S. was supported by the National Science Foundation (DGE 2140743). J.C.W. was supported by NIH grant HL098316. He is an American Cancer Society Research Professor and a member of the Howard Hughes Medical Institute.

## Author Contributions

E.W.S. wrote all code and performed all experiments. E.W.S. and J.C.W. conceived the experiments and wrote the paper.

## Declaration of AI-assisted technologies

Open AI’s GPT-4 was used to assist with coding tasks.

## Methods

### AlphaFold-Multimer (AF-M)

We used a locally installed version of ColabFold^69^ v 1.5.2 to run AF-M. All our predictions used AF-M multimer version 3 weights models 1, 2, and 4 with 3 recycles, templates enabled, 1 ensemble, no dropout, and no AMBER relaxation. The Multiple Sequence Alignments (MSAs) supplied to AF-M were generated by the MMSeq2^70^ server using default settings. All MSAs consisted of vertically concatenated paired and un-unpaired MSAs for the query proteins. The majority of predictions were run on 40GB A100 NVIDIA GPUs while a subset was run on L40S NVIDIA GPUs. Given the memory limitations of these GPUs, we generally cap all jobs at 3,600 amino acids total. In certain cases where a structure is of particular interest, we make exceptions and run AF-M on sequences exceeding 3,600 amino acids.

### Finding contacts in AF-M structures

To determine which residue pairs make valid interfacial contacts in AF-M structures, we use a multi-tiered filtering approach that considers distance along with the pLDDTs and pAE scores. The first step is to iterate through the structure and find all residue pairs in which at least 1 pair of heavy atoms are < 5Å apart and where both residues have pLDDTs > 50. To avoid clashes, we then eliminate pairs with any heavy atoms closer than 1Å. Next, the two PAE scores associated with each residue pair (x, y and y, x) are examined, and if they are < 15 Å, this residue pair is added to the list of valid interfacial contacts. This pipeline differs from our previous approach^50^ where our distance limit was < 8 Å and we did not consider clashes.

### Plots

All plot visualizations were generated using the Python library matplotlib and seaborne running online in Google Colab Jupyter Notebooks.

### PDB human pair contact identification

The PDB API was used to find all cryo-EM and X-ray crystallography structures with overall resolutions < 3.5 Å containing at least two annotated human protein chains. Once all structures with these criteria were retrieved, we iterated through all possible combinations of human chain pairs to find those in contact. Chains were considered to be in contact if they had at least 10 residue pairs with heavy atoms closer than 5 Å. The PDB SIFTS API was used to map PDB chain entries to their corresponding UniProt identifiers.

### Fetching protein sequences

Unless otherwise stated, all protein sequences used for a particular protein represent the full-length isoform sequence reported by the UniProt database at the time of sequence retrieval. To make sequences compatible with AF-M, any non-canonical amino acids not among the standard 20 were removed.

### Random protein sampling

We randomly sampled proteins by randomly shuffling a list of UniProt IDs from the reviewed canonical human proteins downloaded from UniProt and taking the top N ids from this list.

### Extracting and consolidating cross links from publications

We found 20 publications that performed large-scale cross-linking mass spectrometry (XLMS) studies on the scale of whole proteomes or organelles and provided an easily accessible table of identified crosslinks (Table S2). In cases where cross-links were in mouse proteins, we used the UniProt ID mapping tool^71^ to map mouse UniProt IDs to gene symbols to canonical human UniProt IDs. The mouse and human sequences were aligned using the BioPython Align package, and the crosslinked residue numbers were mapped from mouse coordinates to human coordinates. We discarded any crosslinks where this mapping process failed on the ID or residue level. After collecting all unique residue cross-linking pairs, they were deduplicated first on the level of residues and then on the level of protein pairs. To avoid bias, we performed random trimming of over-represented proteins and protein classes. This was done by iteratively identifying the most-common protein across all pairs, randomly selecting all but 28 pairs containing that protein, and then removing them from the list. This process was repeated until no protein was represented more than 28 times across all pairs. After this process, all histone proteins were identified using a list of histone identifiers and randomly removed until histone containing pairs represented only 1% of the final XLMS dataset.

### Random forest training and testing

We used the RandomForestClassifier package from the scikit-learn for our random forest models. Unless otherwise stated, we trained a random forest using criterion=“gini” for split criteria, with 1000 estimators (trees), bootstrapping disabled, and min_samples_split=5. Otherwise, we used the default values specified by scikit-learn. Data was randomly split, with 75% of pairs from each reference set selected for training while the remaining 25% were used for testing. ROC curve visualizations and AUCs were generated via the roc_curve, auc functions imported from the sklearn.metrics package. To generate FDR recall plots, we randomly sub-sampled from the held back test data and constructed test sets with specific ratios of negative to positive examples ranging from ratios of 1:1 to 1:128. In the 1:1 case this corresponded to 403 TP to 403 Tn test pairs while in the 1:128 case, 26 TP and 3,291 TN pairs were used. We tried several iterations of hypermeter optimizations with the Random Forest (number of trees, depth, sample split), but found little to no changes in the performance as measured by AUC.

For the purpose of the online SPOC tool, we developed a smaller model compact SPOC or cSPOC that relies on fewer features and 500 trees. Specifically, we removed all MSA derived features (msa_depth_diff, conservation_mean_diff, perturb_seq_score, conservation_mean, mutual_information_mean; see Table S3) to obviate the need for users to supply MSA files associated with their AF-M runs. Due to their low importance scores, sever features (perturb_seq_score, best_num_phospho_residues, best_num_direct_phospho_contacts, best_num_close_cross_chain_phosphates) were also removed.

### AlphaMissense data processing

Data was downloaded from the human proteome-wide precomputed amino acid substitution data table hosted at (https://console.cloud.google.com/storage/browser/dm_alphamissense;tab=objects?pli=1&prefix=&forceOnObjectsSortingFiltering=false). For each residue position, we averaged the AlphaMissense score across all 19 possible missense variants predicted to produce a single, per-residue missense score. These values were then loaded into a JSON dictionary where each key is a UniProt ID that points to a numeric vector (with the same length as the protein) where each entry represents the averaged missense score for the residue at the corresponding position.

### RNA Coexpression data

mRNA co-expression data for human proteins was downloaded from the online web repository coexpressDB (https://zenodo.org/record/6861444/files/Hsa-u.v22-05.G16651-S245698.combat_pca.subagging.z.d.zip). This download returns a folder with files for each gene such that the name of the file is the ENTREZ gene ID. For every protein/gene we sorted by score (high to low) and took the top 500 pairs before mapping. ENTREZ gene IDs were then mapped to canonical UniProt entry names using the UniProt mapping tool. Pairs where the mapping process failed were discarded.

Score values were used as supplied by the database.

### DEPMAP data

CRISPR KO gene effect data was downloaded from the online resource DEPMAP (https://depmap.org/portal/download/custom/). Every protein was converted into a DEPMAP vector where every entry/dimension corresponds to the Chronos output (gene effect) for that gene in a cell line.

### PerturbSeq data

PerturbSeq mRNA profiles were downloaded from the Figshare data repository (https://plus.figshare.com/articles/dataset/_Mapping_information-rich_genotype-phenotype_landscapes_with_genome-scale_Perturb-seq_Replogle_et_al_2022_processed_Perturb-seq_datasets/20029387). Pearson correlation was then calculated using the numpy package between all combinations of mRNA profiles across all proteins in the dataset. For each protein, the correlation scores were sorted from high to low and the top 200 correlations for every protein were exported.

### Phosphosite data

Phosphorylation data across all organisms was downloaded from the online dbPTM^72^ repository (https://awi.cuhk.edu.cn/dbPTM/download.php). We then extracted the lines matching canonical human UniProt identifiers where each line represents a protein and a corresponding residue location (1 indexed) where a phosphorylation was detected.

### BioGRID ORCS data processing

CRISPR KO data for studies conducted in human cell lines was downloaded from the BioGRID file repository (https://downloads.thebiogrid.org/File/BioGRID-ORCS/Release-Archive/BIOGRID-ORCS-1.1.15/BIOGRID-ORCS-ALL-homo_sapiens-1.1.15.screens.tar.gz). Every gene was mapped to a canonical UniProt ID and for each gene its appearance in the CRISPR screens was converted into binary vectors where each index represents whether that gene was considered a “hit” (0 = no hit, 1 = hit) by the criterion employed by that screen.

### DeepLoc2 protein localization predictions

To have uniform localization information for proteins that went beyond standard and incomplete annotations, we utilized predictions from the DeepLoc 2.0 protein sequence transformer model. We downloaded (https://services.healthtech.dtu.dk/cgi-bin/sw_request?software=deeploc&version=2.0&packageversion=2.0&platform=All) and installed a local copy of DeepLoc 2.0. After installation, we inputted a FASTA file containing all canonical Swiss Prot reviewed sequences for the human proteome downloaded from UniProt. We then used the ESDM1B “fast” model to predict individual localization probabilities split across 10 different possible categories for all sequences. These values were then loaded and stored in a JSON dictionary where each key is a UniProt ID that points to a numeric vector with the 10 localization probabilities output by DeepLoc 2.0.

### H5 protein embeddings

Per-protein embeddings (vectors of length 1024) were retrieved for all reviewed UniProtKB Swiss-Prot human entries via download from UniProt (https://ftp.UniProt.org/pub/databases/UniProt/current_release/knowledgebase/embeddings/UP000005640_9606/per-protein.h5).

### STRINGDB scores

All human protein association scores were downloaded from the STRINGv12 database at (https://stringdb-downloads.org/download/protein.links.detailed.v12.0/9606.protein.links.detailed.v12.0.txt.gz). Each entry in the file lists a pair of proteins identified by their STRINGDB ID consisting of the taxon ID (9606 for humans) concatenated with an ENSEMBL protein id. These ENSEMBL protein ids were mapped to UniProt IDs using UniProt’s mapping API. In cases where this mapping yielded non-canonical UniProt IDs or non-SwissProt entries, these ENSEMBL protein ids were mapped to genes and then each gene was mapped to the canonical Swiss Prot UniProt ID.

### Replisome structural model

An AF-M prediction of the pol ε holoenzyme (POLE1-POLE2-POLE3-POLE4; only POLE1 shown) was aligned on the C-terminal, non-catalytic lobe of POLE1 in a cryo-EM replisome structure (PDB: 7PLO). To model the primer template, the structure of yeast POLE1 catalytic domain with a primer template (PDB: 4M8O) was aligned to the catalytic domain of the AF-M POLE1 structure prediction from the pol ε holoenzyme above. The catalytic domain of POLE1(residues 1-1180), CTF18, CTF8, DCC1 were folded, in which POLE1 made extensive contacts with the CTF18-CTF8-DCC1 complex. The resulting structure was also aligned on POLE1 of the pol ε holoenzyme. Separately, MCM3, MCM7, and CTF18 (only well-ordered residues 281-865) were folded together. MCM3 and the N-terminal lobe of MCM7 (residues 1-319) were deleted, and the remaining C-terminal lobe of MCM7 and CTF18 were aligned on MCM7 from 7PLO.

---

**Figure S1:**
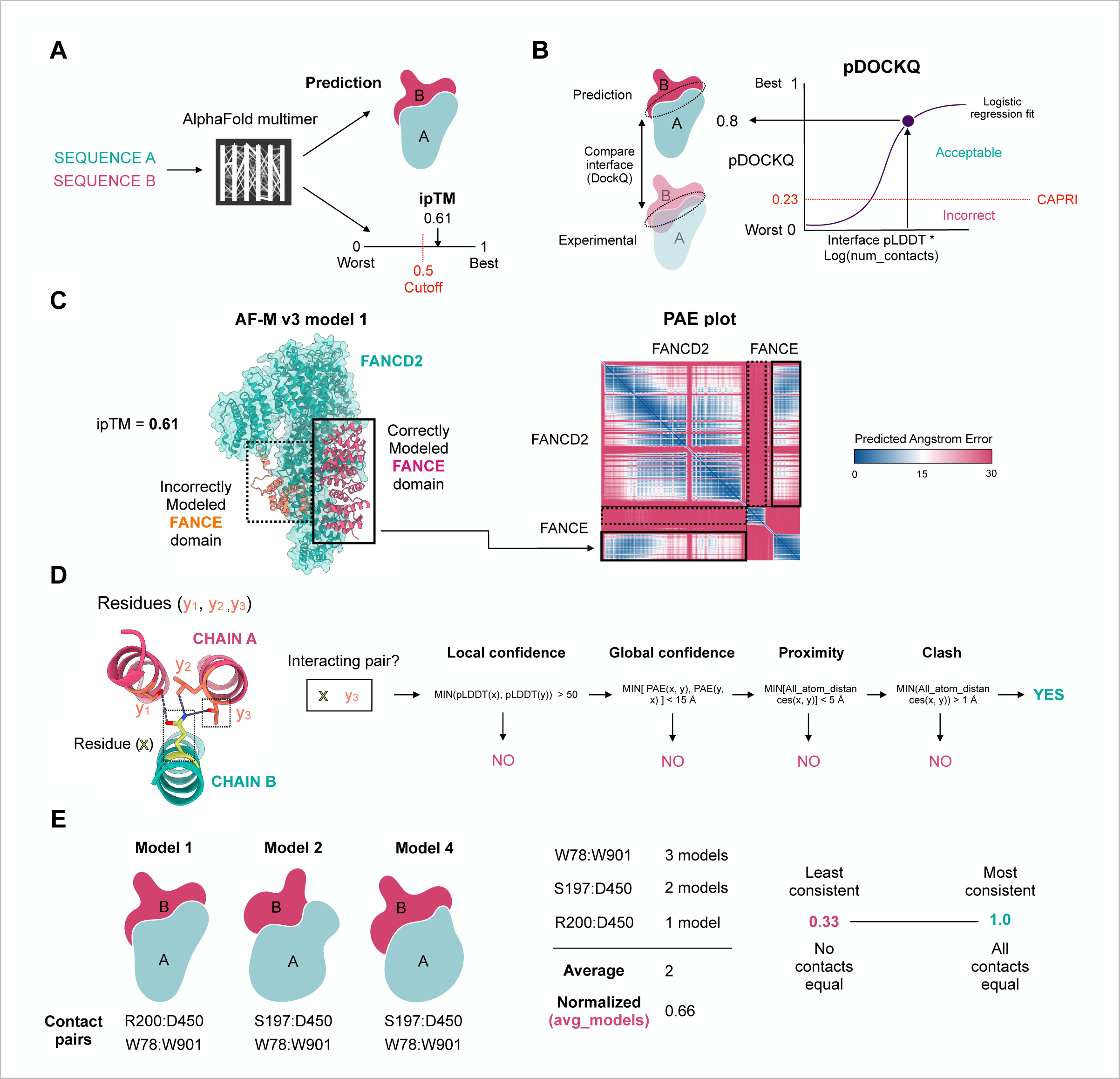
Measuring predicted AF-M interface quality. **(A)** The interface <u>p</u>redicted <u>T</u>emplate <u>M</u>odeling score (ipTM) is a learned AF-M output that estimates how well the predicted interface would match the experimental-determined structure. This score ranges from 0 to 1 (best). A score of > 0.5 is considered a good interaction. **(B)** pDOCKQ is the predicted DOCKQ (docking quotient) score that measures how well a predicted interface matches a ground truth interface between 2 proteins. While DOCKQ can only be calculated in cases where a predicted pair has a corresponding experimental structure, the pDOCKQ can be calculated by passing the average interface pLDDT and number of contacts into a logistic regression formula which was derived by comparing AlphaFold predictions to experimental structures^30^. DOCKQ and pDOCKQ above 0.23 are considered an acceptable match by the CAPRI criteria. **(C)** Considering interface PAE is crucial for interpreting interface quality because AF-M sometimes juxtaposes two well-folded regions with high pLDDTs from two proteins, even if it is not confident about their interaction. In the example shown, proximity (and pDOCKQ) suggests that two regions of FANCE interact with FANCD2. Only by considering the PAE scores of the interfacial residues is it evident that interaction of the C-terminal (red) but not the N-terminal (orange) domain of FANCE with FANCD2 is likely correct. **(D)** All our analysis pipelines first identify residues between chains that satisfy the illustrated criteria. Such “C+” pairs are considered valid interfacial contacts and are used for downstream calculations of interface pLDDT, PAE, etc. **(E)** To quantify the agreement among multiple AF-M predictions of a binary complex, we first identify all valid (C+) residue pairs across found in all the predictions and then count how many predictions each unique contact appears in. This count is averaged across all unique, C+ contacts and then normalized by dividing by the number of predicted structures, yielding the “avg_models” score. In the example shown, one contact is common to all three predictions (W78:W901), one is common to two (S197:D450), and one is unique to one pair (R200:D450), yielding avg_models = (3 + 2 + 1)/ 3 * 3 = 0.66. If all three contacts appeared in all three predictions, avg_models would be 1, indicating perfect agreement between all models.

**Figure S2:**
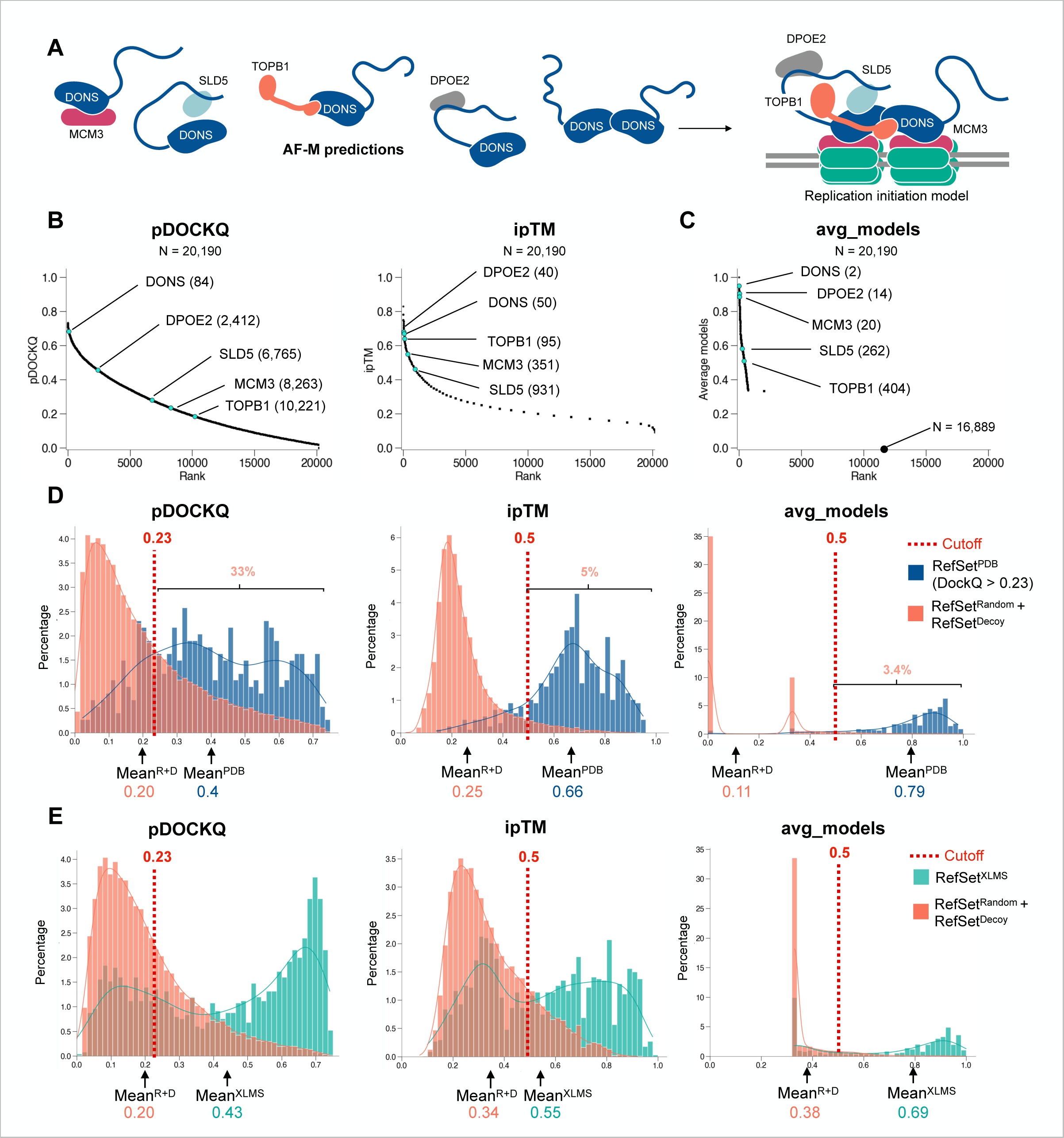
Conventional AF-M confidence metrics have limitations. **(A)** A prior screen for DONSON interactors identified 5 proteins (DPOE2, SLD5, TOPB1, and MCM3, and DONS itself) that were shown to be functional DONSON partners. These binary predictions supported a new model of CMG assembly by DONSON. **(B)** We previously used AF-M to screen DONSON against the entire human proteome and ranked the hits using existing confidence metrics^50^. Here, we display the same data (but recalculated using our updated contact criteria; see methods) as rank plots, which show that pDOCKQ, ipTM, and avg_models scatter DONSON’s interactors over the top ∼1,000-10,000 hits. **(C)** Same as (B) but DONSON hits were ranked by avg_models. **(D)** Distribution of true negative (RefSet^Random^ + RefSet^Decoy^) and true positive (RefSet^PDB^) pairs sorted by pDOCKQ, ipTM, and the avg_models scores. In contrast to other analysis, here all pairs were considered regardless of whether they were contact positive (C+), so that we could observe the performance of these metrics as they are normally applied, independently of our contact criteria. Percentage of pairs exceeding the conventional threshold criteria (dotted line) for each metric is shown above the bracket. **(E)** Distribution of true negative (RefSet^Random^ + RefSet^Decoy^) and training (RefSet^XLMS^) pairs (all C+) sorted by pDOCKQ, ipTM, and the avg_models scores.

**Figure S3:**
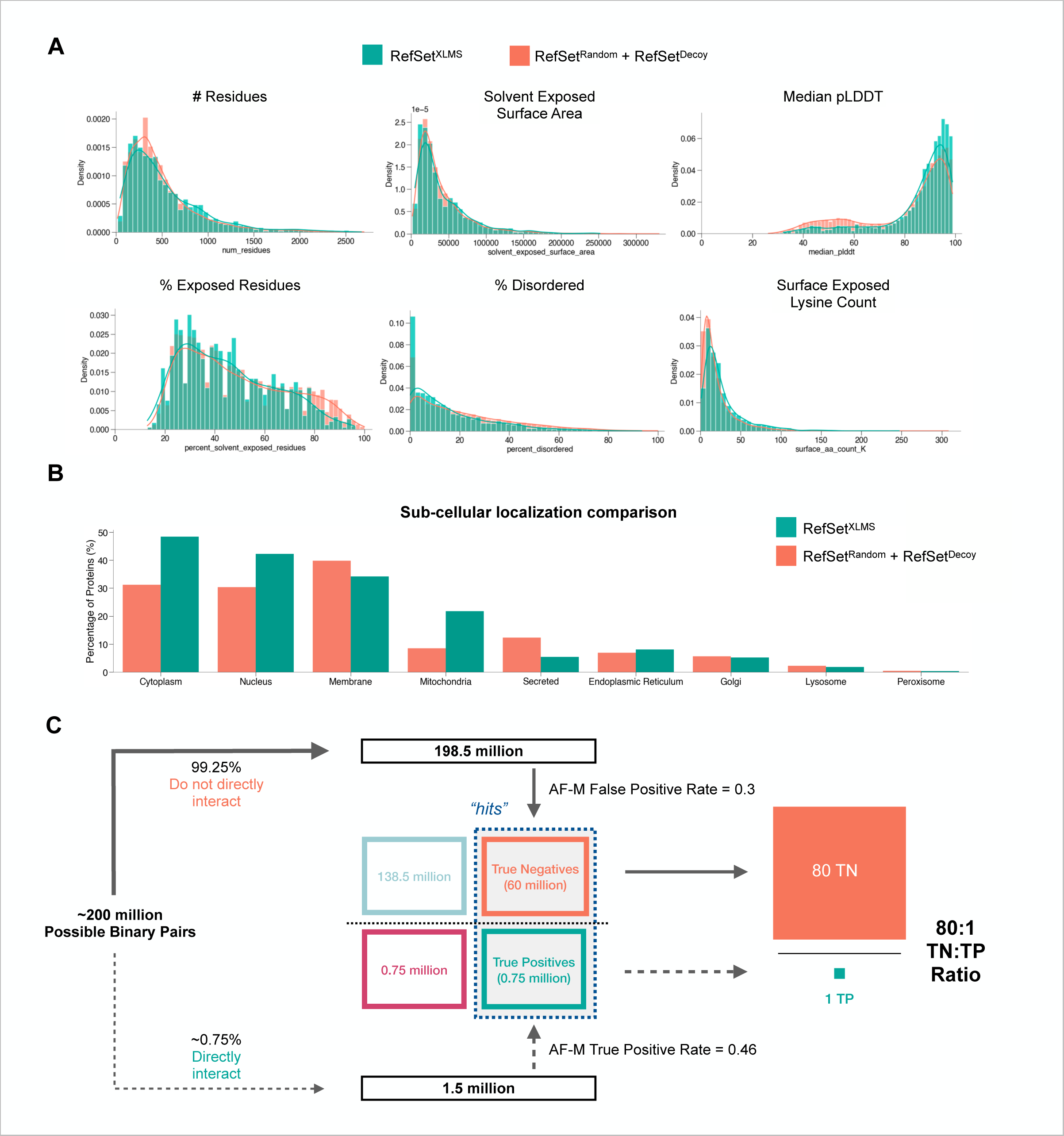
Comparing positive and negative dataset protein composition. **(A)** For all unique proteins in the TN (RefSet^Random^ + RefSet^Decoy^) and training (RefSet^XLMS^) datasets, we downloaded their AlphaFold structures^75^ and calculated various parameters. We verified that RefSet^XLMS^ was not aberrantly enriched for surface-exposed lysines. **(B)** A comparison of proteins in (RefSet^Random^ + RefSet^Decoy^) and RefSet^XLMS^ based on UniProt-annotated subcellular localization. **(C)** Schematic illustrating how we converted the TN:TP ratio in the human binary interactome into the TN:TP ratio that would be observed in an unbiased AF-M screen. We assume 1.5 million TPs and 198.5 million TNs in the human proteome^2^. Based on an AF-M rate of 46.1% in identifying true positives (Figure 2D and accompanying results section), the 1.5 million proteomic TPs would produce 750,000 “hits.” Analogously, because only 30% of random (TN) pairs satisfy the contact (C+) criterion (Figure 2A), 198.5 million TNs would yield ∼60 million contact positive (C+) TN pairs. Dividing 60 million by 750,000 yields an apparent TN:TP ratio of 80:1.

**Figure S4:**
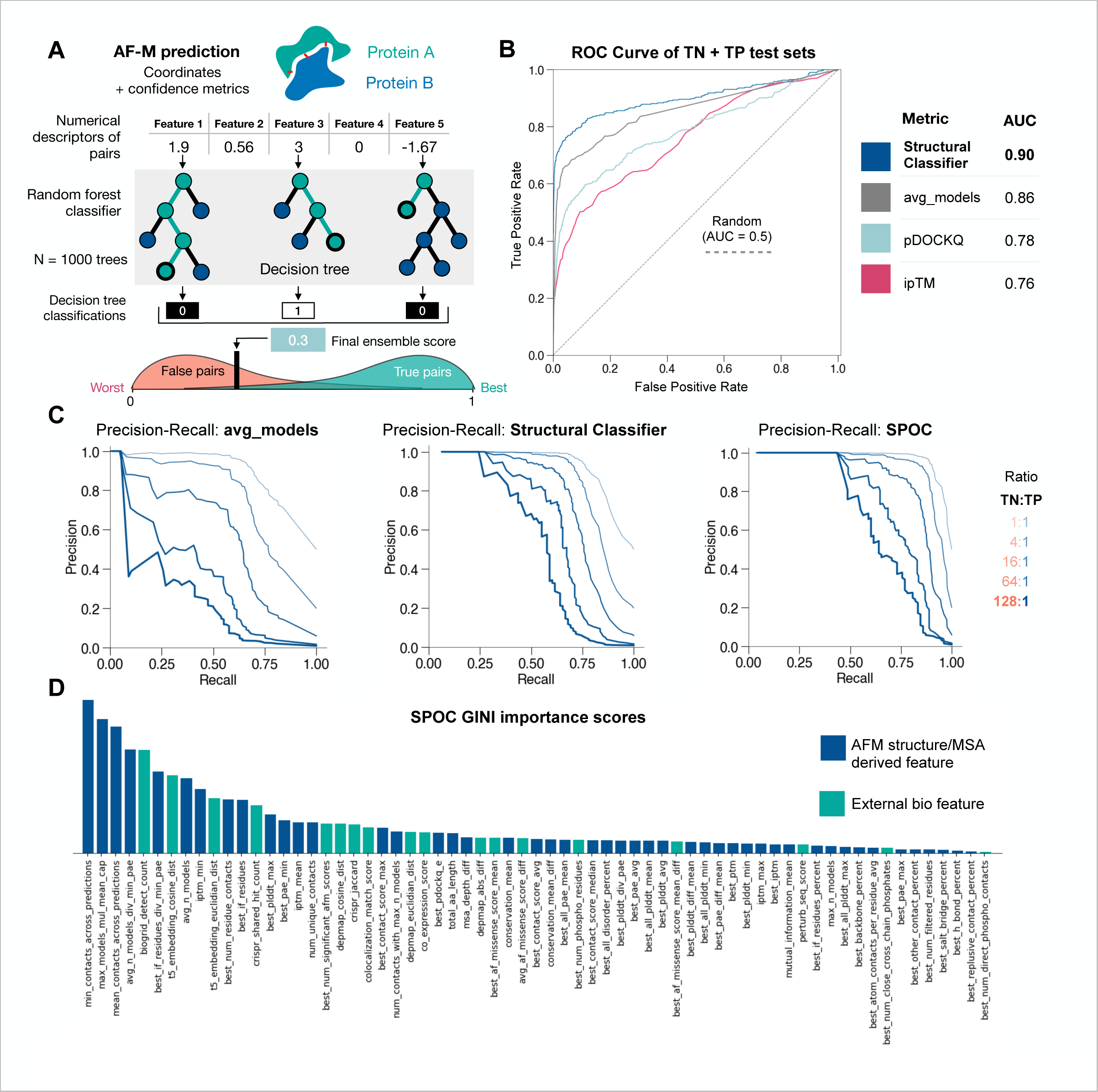
The SPOC classifier for assessing binary predictions. **(A)** Schematic illustrating how an AF-M prediction is converted into a numeric feature representation that can be fed into a random forest machine learning classifier. A random forest is trained by giving it example data with features along with the desired label/class. During training, the forest builds many independent decision trees out of random feature subsets, with each tree attempting to assign the correct label (0 or 1) to the data. Once trained, new instances to be predicted are passed through all the trees, with each tree routing the pair through all its decision layers until it hits a terminal node, where it is assigned a 0 or 1. This final binary vote is tallied across all trees and averaged to produce a final ensemble score. When working effectively, the classifier produces low scores for false pairs and higher scores for true pairs, but for complex data, this separation is generally not perfect. **(B)** AUC (Area Under the Curve) values for ROC (Receiver Operating Characteristic) curves for the structural classifier and other metrics. TN is (RefSet^Random^ + RefSet^Decoy^); TP set is RefSet^XLMS^. The ROC curve shows the True Positive Rate (y) and False positive rate (x) as a function of selected threshold (which is not explicitly show3n on the graph). The stringency increases (higher threshold) moving from the top right to the bottom left of the graph. The better a discriminator can separate false examples from true examples, the more its ROC curve shifts to the upper left, resulting in higher AUC values. **(C)** Precision recall (PR) plots show how recall TP/(TP+FN) and precision TP/(TP+FP) changes as a function of threshold. The best discriminators produce PR curves that tend toward the upper right area of the plot as they recover a high percentage of true examples without including too many false examples. In the plots shown here, SPOC exhibits the best performance, as measured by its curves staying closest to the upper right even as the TN:TP ratio in the test data increases. TN is (RefSet^Random^ + RefSet^Decoy^); TP set is RefSet^XLMS^. **(D)** Histogram of GINI importances. Training a random forest involves constructing decision trees to achieve optimal data splits. All trees record the features they used and how useful they were for sorting data into the correct categories. Splitting usefulness is quantified for every feature via a metric called GINI importance. A higher score indicates that a feature is more helpful in guiding predictions. Plotting GINI importances for SPOC reveals that many features drove its performance.

**Figure S5:**
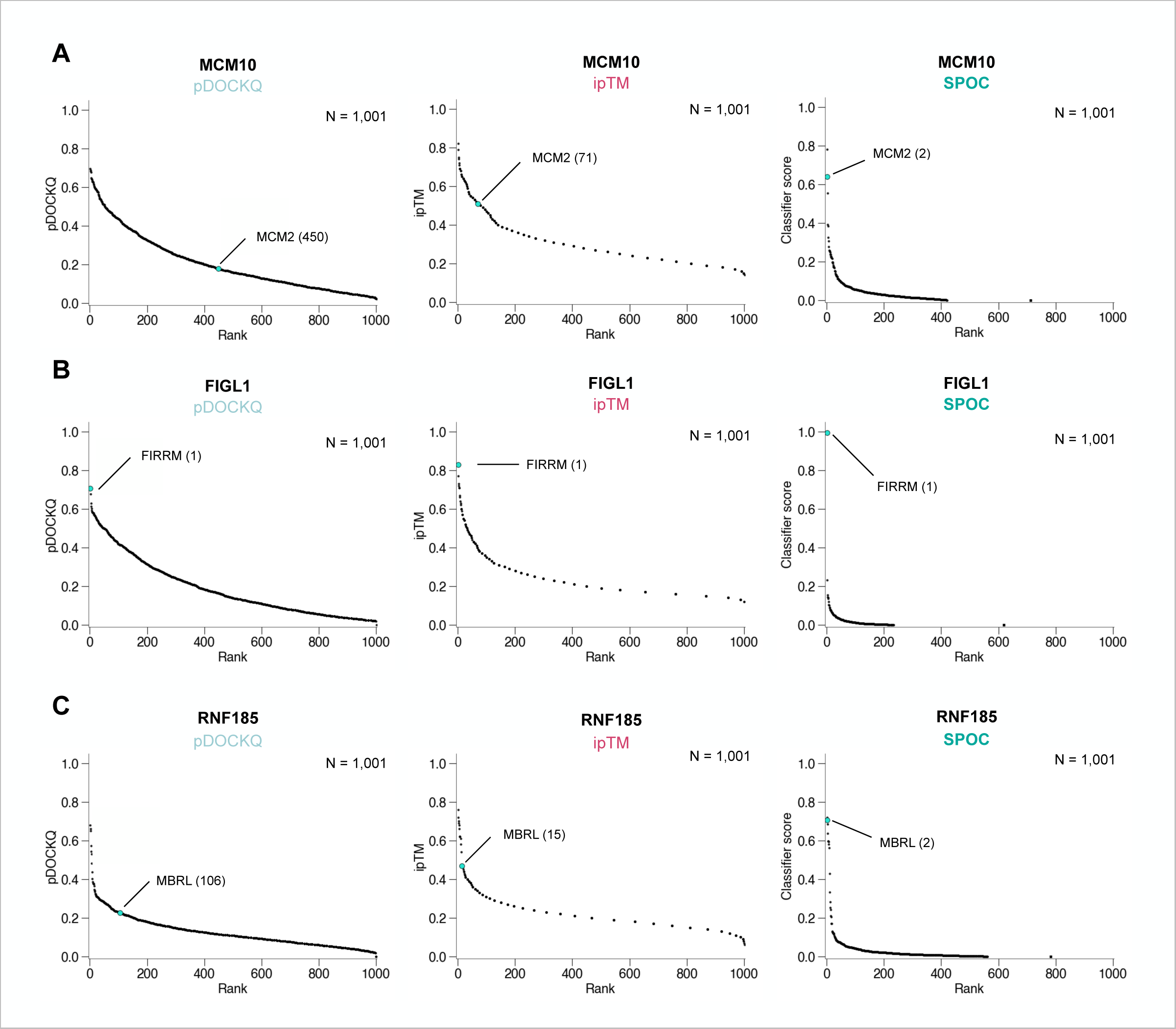
Comparing ranking performance of different metrics. **(A)** The MCM10/MCM2 true pair was embedded in 1000 random pairs involving MCM10, and the AF-M predictions for each pair were ranked using three different metrics. **(B)** The FIGL1-FIRRM true pair was embedded in 1000 random pairs involving FIGL1, and the AF-M predictions for each pair were ranked using three different metrics. **(C)** The RNF185-MBRL true pair was embedded in 1000 random pairs involving RNF185, and the AF-M predictions for each pair were ranked using three different metrics.

**Figure S6:**
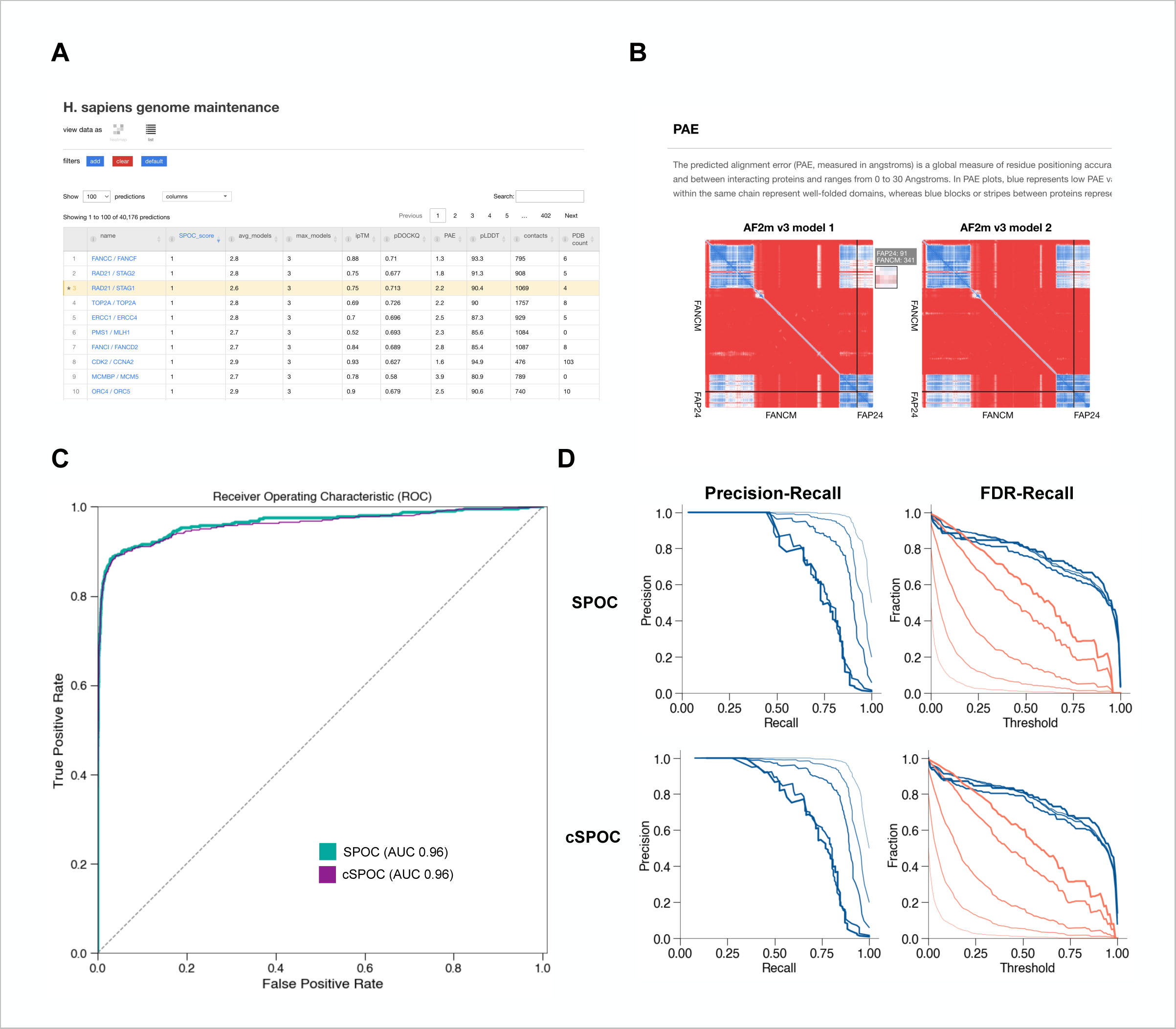
A web portal for viewing and analyzing AF-M predictions. **(A)** Typing a protein name into the search field on the matrix page of predictomes.org retrieves an interactive list of a protein’s hits prioritized by SPOC score. Users can highlight specific rows for visual reference and retrieval. Clicking a row routes the user to the protein’s information page. (**B**) Interactive PAE plots that describe the relative distance error (Å) between any two residues in the predicted complex. Hovering on the plots brings up a smaller view that displays the residue coordinates as well as a closer view of the PAEs near the area of interest. **(C)** ROC plot that compares SPOC to “cSPOC,” which is used to score user-generated AF-M predictions at predictomes.org. Both SPOC and cSPOC have AUCs of 0.96 when benchmarked on the same test data. TN is (RefSet^Random^ + RefSet^Decoy^); TP set is RefSet^XLMS^. **(D)** Precision-recall plots and FDR-recall plots confirm that cSPOC exhibits similar performance as SPOC, even at the highest TN:TP ratios. TN is (RefSet^Random^ + RefSet^Decoy^); TP set is RefSet^XLMS^.

**Figure S7:**
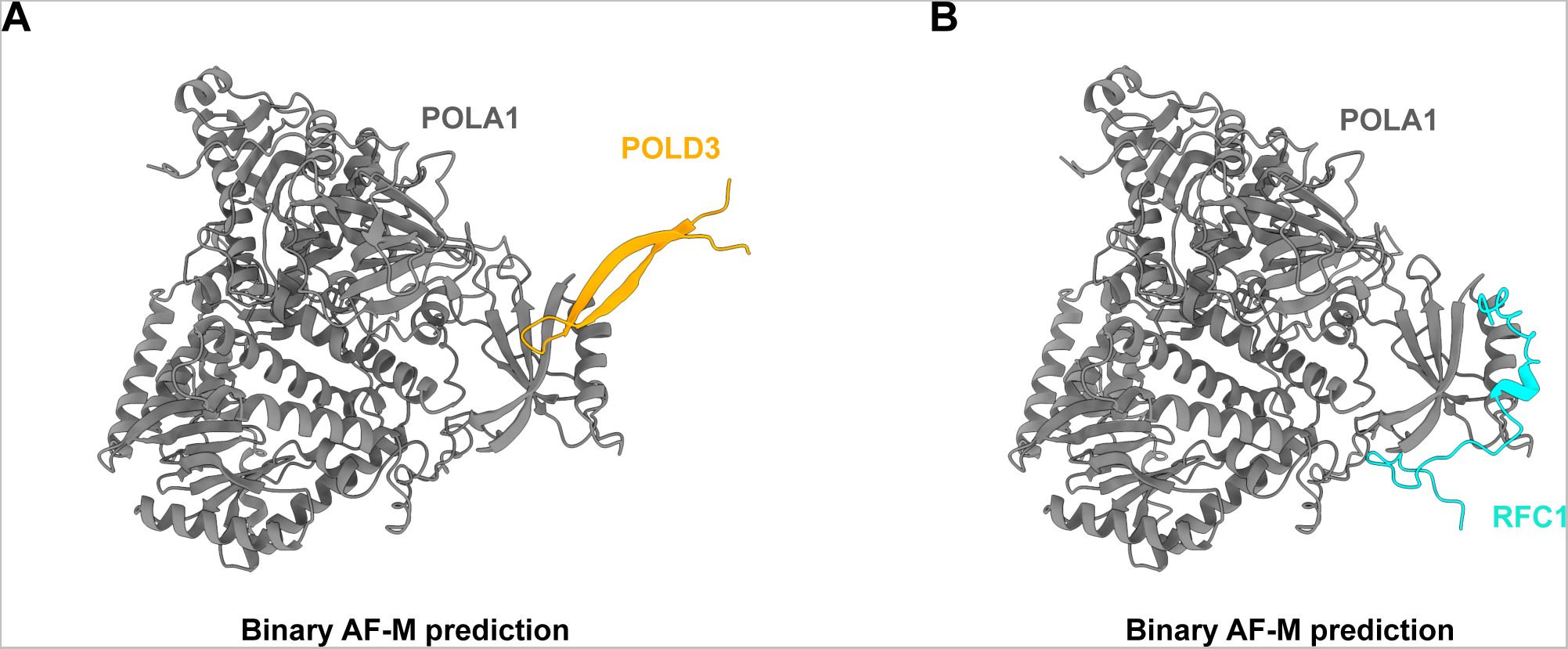
Binary AF-M predictions for Figure 7 model. **(A)** AF-M prediction of the human POLA1-POLD3 complex. The orientation of POLA1 is exactly as in Figure 7B. See Table S5 for confidence metrics. **(B)** AF-M prediction of the human POLA1-RFC1 complex. The orientation of POLA1 is exactly as in Figure 7B. The interaction between RFC1 and POLA1 is less extensive than in the presence of POLD3, as seen in Figure 7B. See Table S5 for confidence metrics.

